# STING activation drives pericentrosomal positioning of acidic organelles via the VAIL-LRRK2-Rab35 pathway

**DOI:** 10.64898/2026.09.11.750829

**Authors:** Shoichi Suenaga, Maria Sakurai, Takeshi Iwatsubo, Tomoki Kuwahara

**Affiliations:** Department of Neuropathology, Graduate School of Medicine, The University of Tokyo, Tokyo, Japan; Department of Dementia Inclusion and Therapeutics, The University of Tokyo Hospital, Tokyo, Japan; National Center of Neurology and Psychiatry, Tokyo, Japan

**Author notes:** Corresponding authors at: 7-3-1, Hongo, Bunkyo-ku, Tokyo, 113-0033, Japan. E-mail address (T. K.), (T. I.).

**Keywords:** centrosome, LRRK2, lysosome, microglia, Parkinson disease, recycling endosome, STING, VAIL

## Abstract

The cGAS-STING pathway is an innate immune system that responds to cytoplasmic double-stranded DNA, but recent studies have highlighted its noncanonical role in facilitating lysosomal stress responses. Specifically, STING activation has been shown to trigger V-ATPase-ATG16L1-induced LC3 lipidation (VAIL) and to activate LRRK2, a Parkinson disease-associated kinase, but the detailed molecular mechanism and the physiological roles of the activation have remained unclear. Here, we found that STING activation in microglia induces the translocation of LRRK2 to the pericentrosomal area, where lysosomes and recycling endosomes (REs) concentrate. This process was VAIL-dependent, and LRRK2 was found to be localized to the organelles. The translocation of LRRK2 required intact microtubule structures and was mediated by LRRK2 kinase activity, its substrate Rab35 and the motor adaptor proteins including JIP4. All these proteins were concentrated at the pericentrosomal area upon STING stimulation. The hyperactivating G2019S mutation of LRRK2 promoted the pericentrosomal translocation of these proteins. Overall, we propose a novel STING-induced noncanonical pathway that involves LRRK2 to facilitate pericentrosomal organelle positioning.

## Introduction

The cyclic GMP-AMP synthase-stimulator of interferon genes (cGAS-STING) pathway is an innate immune signaling axis that responds to cytosolic double-stranded DNA from pathogens as well as mislocalized self-DNA [1]. While this pathway is protective during infection, sustained or excessive STING signaling can become maladaptive, driving sterile inflammation and tissue injury in multiple disease contexts. In neurodegenerative diseases, mitochondrial or genomic stress has been suggested to increase cytosolic self-DNA signals that induce STING-dependent microglial activation, leading to progressive neuroinflammation and neuronal damage [2].

Well-studied functions of the cGAS-STING pathway include the upregulation of type I interferons and other immunomodulatory molecules [3,4], and the activation of inflammatory responses [5,6]. Besides these functions, several studies have reported a noncanonical function of STING that induces robust conjugation of ATG8 family proteins to single membranes of endolysosomes [7–9]. This phenomenon, which does not necessarily involve intracellular degradation, is now referred to as conjugation of ATG8 to single membranes (CASM) [10,11]. Because CASM upon STING activation requires the E3-like ligase ATG16L1, the phenomenon is more specifically referred to as V-ATPase-ATG16L1-induced LC3 lipidation (VAIL) [7,10], in order to distinguish it from a recently identified type of CASM that employs a distinct E3-like ligase, tectonin β-propeller repeat-containing protein 1 (TECPR1) [12–15].

STING-induced VAIL is thought to depend on the recently reported activity of STING as a proton leakage channel [16,17], which perturbs the pH of the Golgi and post-Golgi compartments and engages the V-ATPase-ATG16L1 axis, the core VAIL machinery. Although VAIL requires the ATG conjugation system proteins for its execution, VAIL differs from macroautophagy in that it does not require the upstream autophagy machinery (the ULK1 complex [18] and the class III PI3K complex [19,20]) but requires the C-terminal WD40 domain of ATG16L1 [20]. ATG16L1 lacking its WD40 domain retains autophagic activity but cannot elicit VAIL [20,21].

Leucine rich repeat kinase 2 (LRRK2) is a large serine/threonine kinase that phosphorylates a subset of Rab small GTPases in cells, including Rab8A, Rab10 and Rab35 [22,23]. Missense mutations in *LRRK2* cause autosomal-dominant familial Parkinson disease (PD) [24,25]. Pathogenic mutations enhance LRRK2 kinase activity [26], and converge on increased phosphorylation of its substrate Rab proteins in cells [22,27], leading to dysregulation of the autophagy-lysosome system [28–30], and impaired vesicle transport [31,32]. Hyperactivation of LRRK2 has also been detected in the central nervous system, peripheral blood mononuclear cells, and urine of idiopathic PD cases [33–36].

Under lysosomal stress, LRRK2 localizes to the stressed endolysosomal membranes [37–39], where it becomes activated, regulates lysosomal morphology, and promotes lysosomal exocytosis [37,40]. Several Rab proteins phosphorylated by LRRK2 associate with kinesin/dynein adaptor proteins, thereby modifying subcellular positioning [41,42] and morphology [39,43] of organelles along microtubules. Recently, LRRK2 has been reported to be activated upon STING activation [8,44], leading to the notion that LRRK2 acts as a plausible downstream effector of STING-induced VAIL. Thus, increased signaling from STING to LRRK2 may contribute to PD.

Despite these advances, detailed intracellular functions of LRRK2 downstream of the cGAS-STING pathway have remained unknown. Because LRRK2 participates in the regulation of stressed endolysosomes via VAIL, we hypothesized that STING activation leads to the recruitment of LRRK2 to V-ATPase-positive organelles and thereby modifies their subcellular distribution. In this study, we discovered that STING activation induces translocation of LRRK2 as well as lysosomes and recycling endosomes (REs) toward the pericentrosomal area. We further demonstrated that this translocation is primarily mediated via the VAIL-LRRK2-Rab35 pathway.

## Results

### LRRK2 is translocated to the pericentrosomal area upon STING activation

Given the possible role of STING-induced VAIL in LRRK2 activation, we first analyzed the detailed changes in the activity and subcellular localization of LRRK2 under STING activation. We chose the MG6 mouse microglia cell line for the analyses because these cells endogenously express a substantial level of LRRK2 when primed by interferon-γ (IFNγ), which induces polarization toward the pro-inflammatory M1 phenotype [45–47]. Consistent with previous reports [44], treatment of MG6 cells with the STING agonist 2′,3′-cyclic GMP-AMP (cGAMP) potently induced phosphorylation of Rab10 at Thr73, an established readout of cellular LRRK2 kinase activity [22,23], and this phosphorylation was abolished by treatment with an irreversible STING inhibitor, H-151 [48] (**Figures 1A,B**). We confirmed the activation of the conventional STING pathway upon cGAMP treatment, as indicated by TBK1 phosphorylation and STING disulfide-linked dimerization [49,50] (**Figures 1A,C-E**). The increase in TBK1 phosphorylation was suppressed by H-151 treatment, while dimerization of STING was not, both of which were consistent with previous reports [51,52]. The activation of LRRK2 was also induced by another STING agonist diABZI (**Fig. S1A-F**), further supporting that LRRK2 is activated downstream of STING.

**Figure 1.**
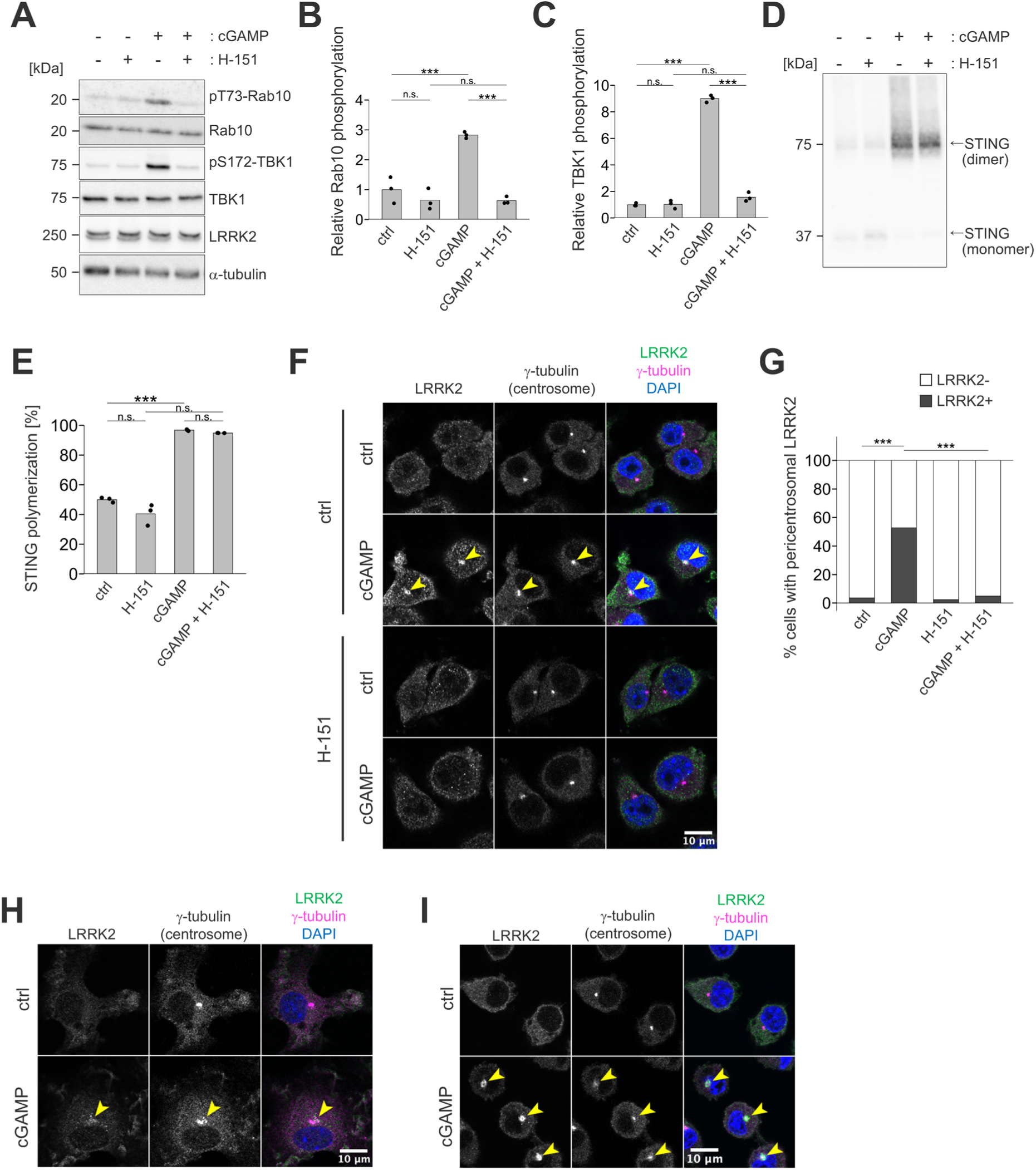
LRRK2 is translocated to the pericentrosomal area upon STING activation. (**A**) Immunoblotting of lysates of MG6 cells primed by IFNγ and treated with cGAMP and/or STING inhibitor H-151 for six hours. Representative images from three independent sample sets. (**B**, **C**) Quantification of Rab10 phosphorylation (B) and TBK1 phosphorylation (C), as shown in A. Statistical significance was assessed by two-way ANOVA with Tukey’s post-hoc test. (**D**) Non-reducing immunoblotting of the same sets of cell lysates as in A probed with an anti-STING antibody. (**E**) Quantification of STING dimerization in D. Statistical significance was assessed by two-tailed Welch’s *t*-tests with Holm-Bonferroni’s correction for multiple comparison. (**F**) Confocal immunocytochemical analysis of cGAMP-treated MG6 cells fixed and stained for LRRK2 and other marker proteins as indicated. Arrowheads indicate the LRRK2-positive centrosomes. (**G**) Quantification of the percentages of cells harboring the LRRK2-positive centrosome in F. Only cells in which the γ-tubulin-positive centrosome was within the focal plane and did not overlap the nuclear signal were scored (see Methods). Statistical significance was assessed by chi-squared tests with Holm-Bonferroni’s correction. (**H**) Immunocytochemical analysis of IFNγ-primed primary microglia treated with cGAMP for six hours, fixed, and stained for LRRK2 and other markers as indicated. Arrowheads indicate LRRK2-positive centrosomes. (**I**) Immunocytochemical analysis of RAW cells primed by IFNγ and treated with cGAMP for six hours. Arrowheads indicate LRRK2-positive centrosomes. n.s.: not significant, ***: *p* < 0.001.

We next examined the intracellular localization of endogenous LRRK2 under STING activation by immunocytochemistry and found that LRRK2 often formed distinctive puncta at the pericentrosomal area (**Figure 1F**). LRRK2 puncta consistently encompassed the γ-tubulin signal, appearing as structures slightly larger than, and centered on, the centrosome. The proportion of cells with pericentrosomal LRRK2 puncta was markedly increased by cGAMP treatment (**Figure 1G**). This LRRK2 translocation to the pericentrosomal area was inhibited by H-151 treatment, verifying STING’s involvement (**Figures 1F,G**). The translocation of LRRK2 was also detected in primary mouse microglia (**Figure 1H**) and in the mouse macrophage RAW264.7 cells (**Figure 1I**). LRRK2-positive pericentrosomal puncta in RAW264.7 cells appeared to be larger compared to those in MG6 cells, and some of the LRRK2 puncta were observed as hollow or ring-like, hinting that LRRK2 localizes not to the centrosome itself but to organelles clustered around it. Such pericentrosomal translocation of LRRK2 was not observed in other cell types (*e.g.*, NIH/3T3, HEK293 cells). In NIH/3T3 cells, cGAMP treatment induced phosphorylation of TBK1 and STING dimerization, indicating successful uptake of cGAMP and intended STING activation, but phosphorylation of Rab10 was not enhanced (**Fig. S2A,B**). cGAMP treatment also did not induce LRRK2 translocation in either NIH/3T3 cells or HEK293 cells transiently overexpressing 3×FLAG-tagged LRRK2 (**Fig. S2C,D**). These observations suggest that pericentrosomal translocation and the resultant activation of LRRK2 occur specifically in microglia and macrophage lineage cells.

Previous studies have shown that overactivation of LRRK2 causes deficits in centriole cohesion [53–55]. We observed some split centrosomes in MG6 cells, but the frequency of centrosome splitting did not differ between cGAMP-treated and untreated conditions (**Fig. S3A,B**). Another consequence of excessive LRRK2 activation around the centrosome is the inhibition of primary cilia formation [23,56–58]. However, microglia are known to lack developed primary cilia [59,60], and indeed, no ciliary structure was detected in MG6 cells regardless of cGAMP treatment (**Fig. S3C**).

### Lysosomes and REs are translocated to the pericentrosomal area together with LRRK2 upon cGAMP stimulation

Because VAIL is initiated by the interaction between the WD40 domain of ATG16L1 and the V-ATPase on acidified single membranes [10,20,61], LRRK2 recruitment under VAIL-inducing stress occurs on V-ATPase-bearing endomembrane compartments. We examined the spatial relationship between LRRK2 and these compartments upon STING activation. MG6 cells treated with cGAMP frequently showed accumulation of LAMP1-positive lysosomes in the pericentrosomal region (**Figure 2A**). Of note, LRRK2 puncta were confined to a narrower region around the centrosome than LAMP1-positive structures (**Figure 2B**), indicating that LRRK2 marks a subset of the lysosomes. We defined the lysosome-centrosome average distance (LCAD) as the mean distance from the centrosome to all pixels within the cytosol, weighted by LAMP1 intensity. This measurement was suitable for the images in this study because lysosomes are densely clustered near the centrosome and cannot be resolved as individual structures. Measurement of LCADs quantitatively showed that the distribution of lysosomes shifted toward the vicinity of the centrosome after cGAMP treatment (**Figure 2C**). Moreover, among cGAMP-treated cells, LRRK2 puncta-positive cells exhibited shorter LCADs compared to LRRK2 puncta-negative cells, suggesting that LRRK2 accumulation is associated with redistribution of lysosomes toward the centrosome (**Figure 2D**).

**Figure 2.**
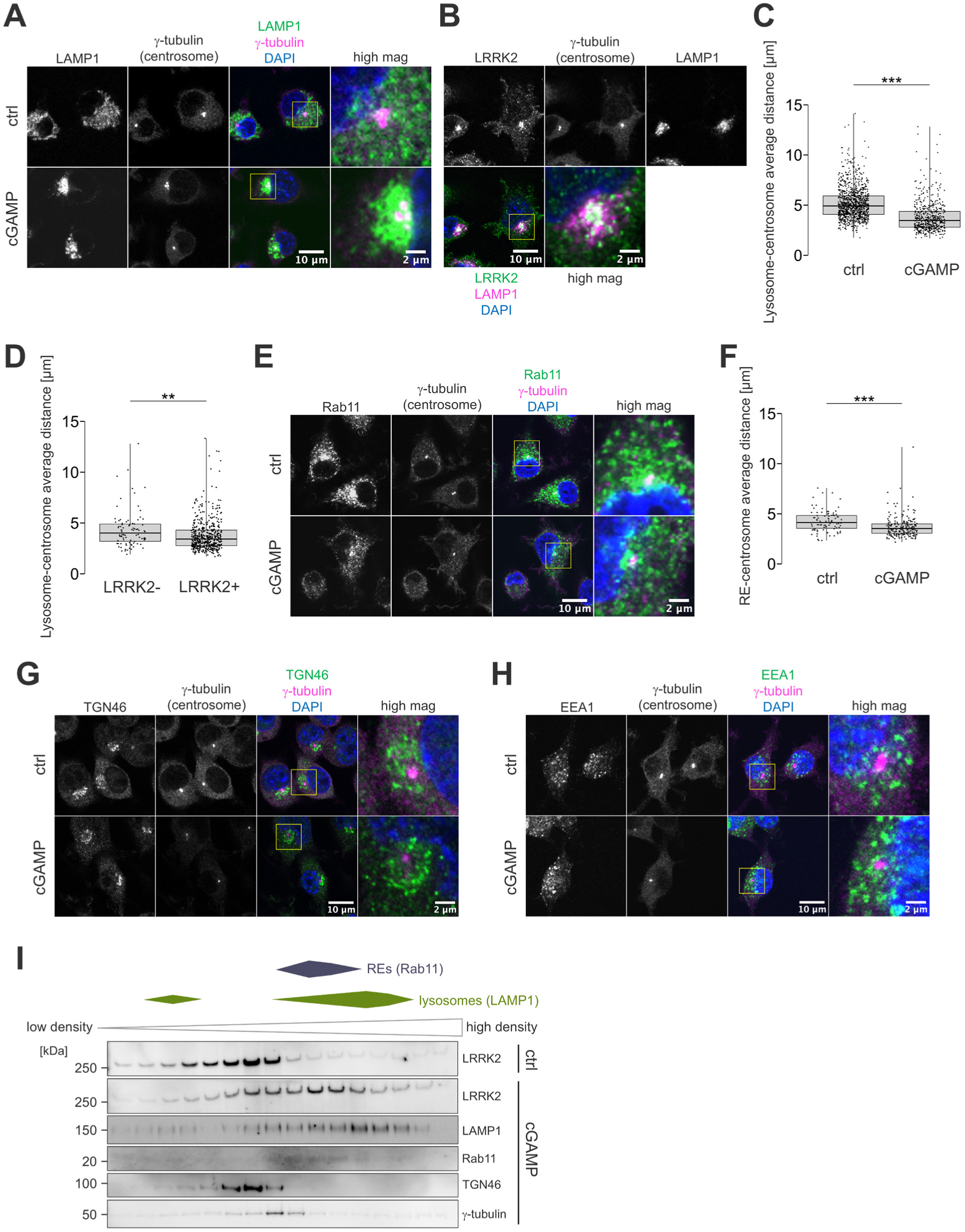
Lysosomes and REs are distributed toward the pericentrosomal area together with LRRK2 upon STING activation. (**A**) Immunocytochemical analysis for LAMP1 and other marker proteins as indicated in MG6 cells treated with or without cGAMP. (**B**) Immunocytochemical analysis showing clustering of LRRK2 and LAMP1 at the pericentrosomal area. (**C**) LCADs in MG6 cells treated with or without cGAMP. Statistical significance was assessed by two-tailed *t*-test. 1060 (control) and 555 (cGAMP-treated) cells were included from two independent experiments. (**D**) LCADs in cGAMP-treated MG6 cells with LRRK2-negative or LRRK2-positive centrosomes. Statistical significance was assessed by two-tailed *t*-test. 90 (LRRK2-) and 517 (LRRK2+) cells were included. (**E**) Immunocytochemical analysis for Rab11 and other marker proteins as indicated. (**F**) RE-centrosome average distances in MG6 cells treated with or without cGAMP. Statistical significance was assessed by two-tailed *t*-test. 87 (control) and 208 (cGAMP-treated) cells were included. (**G**, **H**) Immunocytochemical analysis for TGN46 (G) or EEA1 (H) and other marker proteins as indicated in MG6 cells treated with or without cGAMP. (**I**) Immunoblotting of MG6 homogenates fractionated by sucrose density gradient centrifugation and probed with antibodies for LRRK2 and indicated organelle markers. **: *p* < 0.01, ***: *p* < 0.001.

STING itself is known to be transported to lysosomes for degradation during the signal termination process [3,62,63], but recent studies have highlighted the role of recycling endosomes (REs) as intermediates in the transport of STING to lysosomes [64–66]. We found that, similar to lysosomes, Rab11-positive REs were observed in a punctate pattern at the pericentrosomal area in MG6 cells (**Figure 2E**). Although RE localization near the centrosome was observed even in the absence of cGAMP, it became more pronounced after cGAMP treatment. Quantification of the proximity of REs to the centrosome using the same approach used for lysosomes revealed the shifted distribution of REs toward the centrosome upon STING activation (**Figure 2F**). Together, these observations suggest that lysosomes and REs are translocated to the pericentrosomal area downstream of STING activation.

Because the V-ATPase resides not only on lysosomes or REs but throughout the acidified endomembrane system, we also examined two other V-ATPase-positive compartments that are known to reside near the centrosome: the *trans*-Golgi network and early endosomes. In contrast to LRRK2 puncta, which encompassed the γ-tubulin signal, the *trans*-Golgi marker TGN46 was absent on the γ-tubulin-positive region (**Figure 2G**). The early endosome marker EEA1 also did not overlap with γ-tubulin (**Figure 2H**). Thus, neither compartment occupies the region where LRRK2 puncta form, arguing against the possibility that pericentrosomal LRRK2 is recruited to the *trans*-Golgi or to early endosomes.

Based on the observations on the distribution of lysosomes and REs, we hypothesized that the puncta formation of LRRK2 at the pericentrosomal area is not due to its direct association with centrosomes, but rather represents the recruitment of LRRK2 to V-ATPase-positive organelles that accumulate at the pericentrosomal area. To confirm this hypothesis, we performed organelle fractionation by sucrose density gradient centrifugation. Under unstimulated conditions, LRRK2 was recovered predominantly in low-density fractions, which contained TGN46 but little LAMP1 or Rab11. After cGAMP treatment, an additional peak of LRRK2 appeared in high-density fractions that coincided with the peaks of LAMP1 and Rab11 (**Figure 2I**). Although co-fractionation alone does not establish physical association, this cGAMP-induced redistribution of LRRK2 into lysosome-and RE-containing fractions is consistent with the imaging data above, and together they support the notion that STING activation induces translocation of LRRK2 to lysosomes or REs that accumulate at the pericentrosomal area.

### STING-induced pericentrosomal translocation requires VAIL and GABARAP/GABARAPL1

Since the activation and the lysosomal recruitment of LRRK2 are mediated via VAIL [67] (**Figure 3A**), which is induced by STING activation [7,9,44,68], we next sought to examine whether VAIL machinery is indeed involved in STING-mediated phenomena described above. We first confirmed that cGAMP treatment induced the lipidation of LC3B in MG6 cells, ensuring the execution of VAIL (**Figures 3B,C**). Additionally, we found that VAIL-related ATG8 family proteins LC3B and GABARAP, as well as ATG16L1, were translocated to the pericentrosomal area upon cGAMP treatment with punctate patterns similar to that of LRRK2 (**Figures 3D-G**).

**Figure 3.**
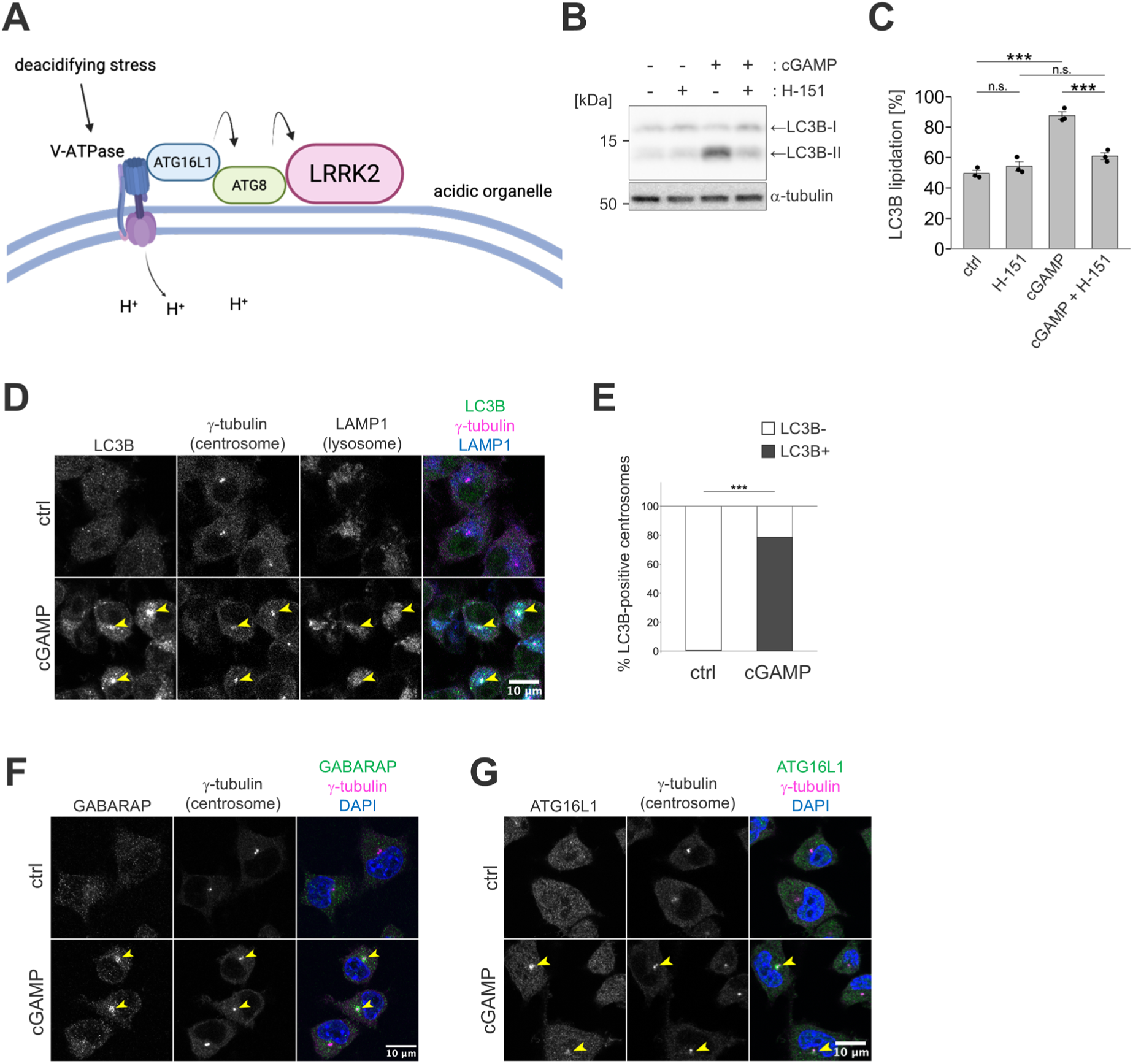
STING activation induces pericentrosomal translocation of VAIL-related molecules. (**A**) A cartoon depicting VAIL-dependent LRRK2 recruitment onto an acidic organelle. Deacidifying stress induces V-ATPase assembly and the subsequent recruitment of ATG16L1 via its WD40 domain, thereby causing the lipidation of ATG8 proteins on the single membrane. LRRK2 is also recruited onto the membrane via VAIL. (**B**) Immunoblotting of lysates of MG6 cells treated with cGAMP and/or H-151 for six hours. Representative images from three independent sample sets. (**C**) Quantification of LC3B lipidation in B. Statistical significance was assessed by two-way ANOVA with Tukey’s post-hoc test. (**D**) Immunocytochemical analysis of cGAMP-treated MG6 cells fixed and stained for LC3B and other markers as indicated. Arrowheads indicate the LC3B-positive centrosomes. (**E**) Quantification of the percentages of cells harboring the LC3B-positive centrosome in D. Statistical significance was assessed by a chi-squared test. Only cells in which the γ-tubulin-positive centrosome was within the focal plane and did not overlap the nuclear signal were scored (see Methods). (**F**, **G**) Immunocytochemical analysis for GABARAP, ATG16L1 and indicated markers in MG6 cells treated with or without cGAMP. Arrowheads indicate the centrosomes positive for GABARAP or ATG16L1.

To examine the involvement of VAIL in the pericentrosomal translocation of LRRK2 and lysosomes, cells were treated with bafilomycin A_1_ (BafA1), a potent V-ATPase inhibitor that suppresses VAIL [11]. BafA1 treatment blocked the cGAMP-induced pericentrosomal translocation of LRRK2 (**Figures 4A,B**). The shortening of LCAD was abolished by treatment with BafA1, suggesting the involvement of VAIL in lysosomal positioning (**Figure 4C**). On the other hand, treatment with saliphenylhalamide (SaliP), a VAIL inducer that promotes association of two subunits of V-ATPase [10], induced LRRK2 translocation to the pericentrosomal area as well as to the enlarged lysosomes in the cytoplasm (**Figure 4D**). We also examined the effect of knockdown of ATG16L1, which associates with V-ATPase V1 subunit and mediates VAIL [20,69]. Since higher knockdown efficiency was achieved in RAW264.7 cells than in MG6 cells, RAW264.7 cells were used in subsequent experiments where knockdown was required. Knocking down endogenous *Atg16l1* blocked the pericentrosomal translocation of LRRK2 upon cGAMP treatment (**Figure 4E,F**). Knockdown efficiency was confirmed in our previous study [70]. These data suggest a crucial role of VAIL in the translocation of LRRK2 under STING activation.

**Figure 4.**
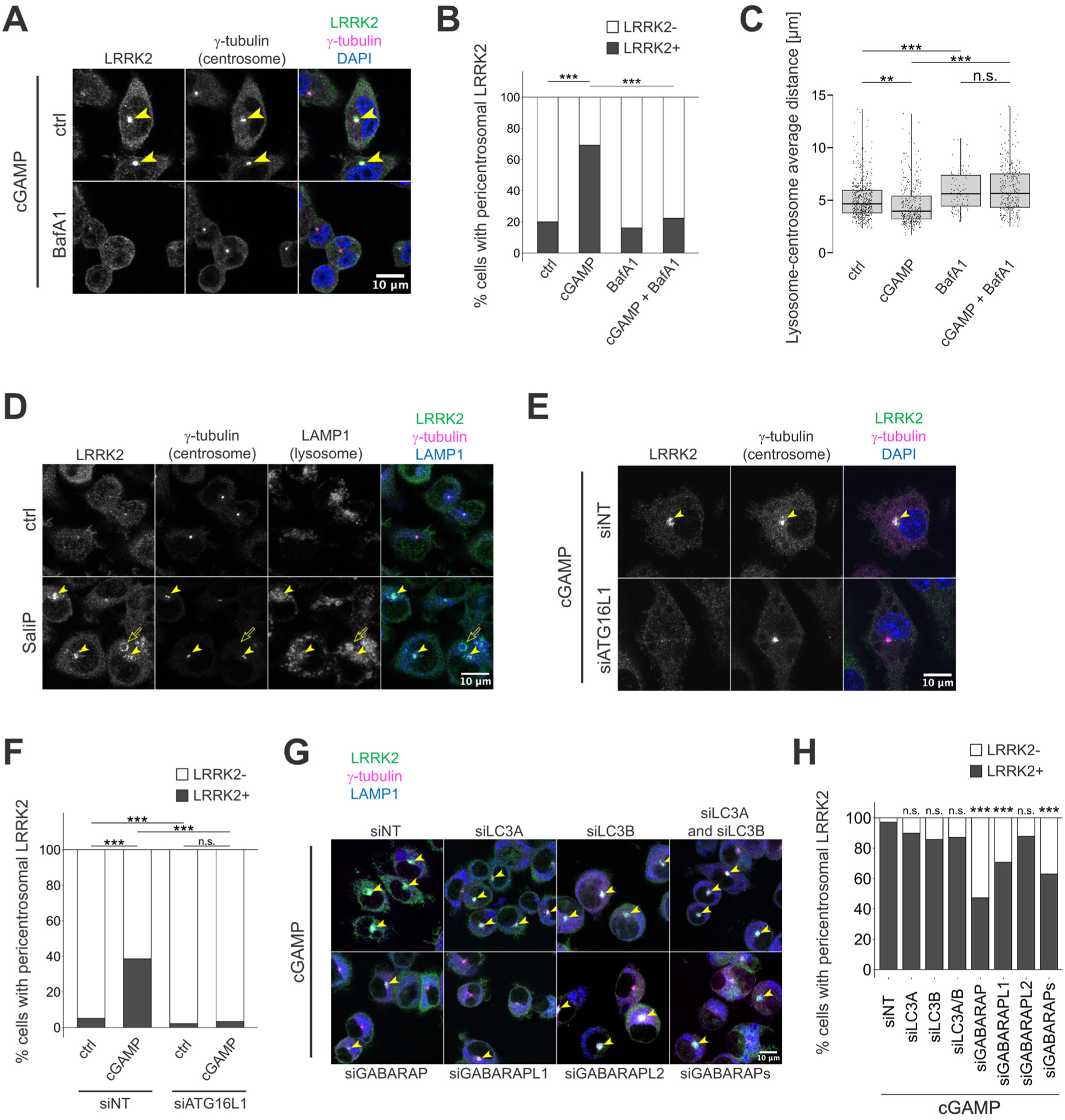
The pericentrosomal translocation of LRRK2 and lysosomes depends on VAIL. (**A**) Immunocytochemical analysis of MG6 cells treated with cGAMP and with or without BafA1. Arrowheads indicate the LRRK2-positive centrosomes. (**B**) Quantification of the percentages of cells harboring the LRRK2-positive centrosome in A, scored as in Fig. 1G. Statistical significance was assessed by chi-squared tests with Holm-Bonferroni’s correction. (**C**) LCADs in MG6 cells treated with cGAMP and/or BafA1. Statistical significance was assessed by two-way ANOVA with Tukey’s post-hoc test. 518, 325, 97, and 305 cells were included in each condition. (**D**) Immunocytochemical analysis of MG6 cells treated with SaliP for three hours. Filled arrowheads indicate the LRRK2-positive centrosomes, and outlined arrows indicate cytosolic large lysosomes. (**E**) Immunocytochemical analysis of RAW264.7 cells knocked down for *Atg16l1*. Arrowheads indicate the LRRK2-puncta. (**F**) Quantification of the percentages of cells harboring the LRRK2-puncta in E. Statistical significance was assessed by chi-squared tests with Holm-Bonferroni’s correction. (**G**, **H**) Immunocytochemical analysis of RAW264.7 cells treated with siRNAs targeting ATG8 family proteins, as indicated. Arrowheads indicate the LRRK2-positive centrosomes. Statistical significance was assessed by chi-squared tests with Holm-Bonferroni’s correction between siNT and each siRNA-treated sample. n.s.: not significant, **: *p* < 0.01, ***: *p* < 0.001.

Next, we sought to identify which members of the ATG8 family recruited by VAIL are involved in the STING-induced LRRK2 translocation. Five major ATG8 family members annotated in the mouse genome were examined: LC3A, LC3B, GABARAP, GABARAPL1, and GABARAPL2. Knockdown of GABARAP or GABARAPL1 in RAW264.7 cells suppressed the frequency of LRRK2 translocation to the pericentrosomal area, whereas knockdown of LC3A, LC3B, or GABARAPL2 did not significantly suppress the frequency in comparison to the control (**Figures 4G,H**). Knockdown efficiencies were confirmed by RT-qPCR (**Fig. S4A-E**). This suggests that GABARAP and GABARAPL1 play a more important role in the pericentrosomal translocation events than LC3s and GABARAPL2.

### The LRRK2-Rab35 axis is responsible for the pericentrosomal translocation of LRRK2

We next tried to explore the molecular mechanism of pericentrosomal translocation downstream of the VAIL-LRRK2 pathway. First, we examined the requirement of LRRK2 kinase activity and found that treatment with the potent LRRK2 kinase inhibitor MLi-2 [71] suppressed the cGAMP-induced LRRK2 translocation to the pericentrosomal area (**Figures 5A,B**). MLi-2 treatment also suppressed cGAMP-induced phosphorylation of Rab10 but not that of TBK1 (**Figures 5C-E**). Furthermore, the shortening of LCAD was abolished upon MLi-2 treatment, indicating that LRRK2 kinase activity is required for the cGAMP-induced repositioning of lysosomes (**Figure 5F**).

**Figure 5.**
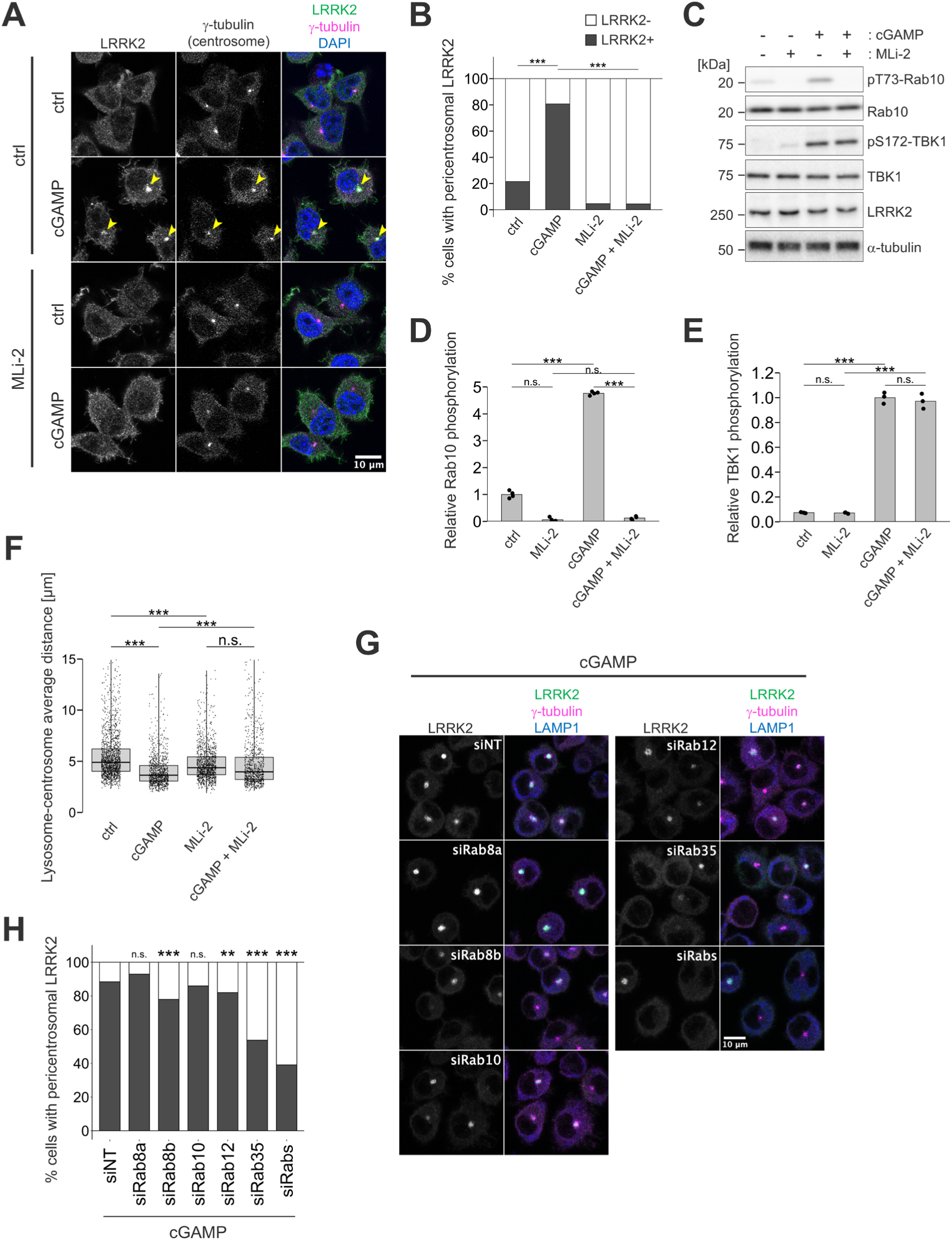
The pericentrosomal translocation of LRRK2 and lysosomes depends on the kinase activity of LRRK2 and its substrate Rab35. (**A**) Immunocytochemical analysis of MG6 cells treated with cGAMP and/or LRRK2 kinase inhibitor MLi-2. Arrowheads indicate the LRRK2-positive centrosomes. (**B**) Quantification of the percentages of cells harboring the LRRK2-positive centrosome in A, scored as in Fig. 1G. Statistical significance was assessed by chi-squared tests with Holm-Bonferroni’s corrections. (**C**-**E**) Immunoblotting of lysates of MG6 cells treated with cGAMP and/or LRRK2 kinase inhibitor MLi-2 for six hours. Representative images from four independent sample sets. Statistical significance was assessed by two-way ANOVAs with Tukey’s post-hoc tests in D and E. (**F**) LCADs of MG6 cells treated with cGAMP and/or MLi-2. Statistical significance was assessed by two-way ANOVA with Tukey’s post-hoc test. 1548, 981, 1205, and 896 cells were included in each condition. (**G**) Confocal immunocytochemistry analysis in RAW264.7 cells knocked down for each Rab protein. (**H**) Quantification of the percentages of cells harboring the LRRK2-positive centrosome in G, scored as in Fig. 1G. Statistical significance was assessed by chi-squared tests with Holm-Bonferroni’s correction between siNT and each siRNA-treated sample. n.s.: not significant, **: *p* < 0.01, ***: *p* < 0.001.

Since LRRK2 kinase activity results in the phosphorylation of a subset of Rab GTPases, we focused on the role of these substrates: Rab8a, Rab8b, Rab10, Rab12, and Rab35 [22,23]. Among these, knockdown of Rab35 most potently suppressed the cGAMP-induced puncta formation of LRRK2 in RAW264.7 cells (**Figures 5G,H**). Knockdown of other Rabs had weaker or little effects. Knockdown efficiencies were confirmed in our previous studies [70,72].

Along with this, immunocytochemical analysis revealed that Rab35 was recruited to the pericentrosomal area after cGAMP treatment (**Figures 6A,B**), suggesting a role for Rab35 downstream of LRRK2 under cGAS-STING activation. Of note, other phosphorylated Rabs including Rab8 and Rab10 were also enriched at the pericentrosomal area upon STING activation (**Figures 6C,D**).

**Figure 6.**
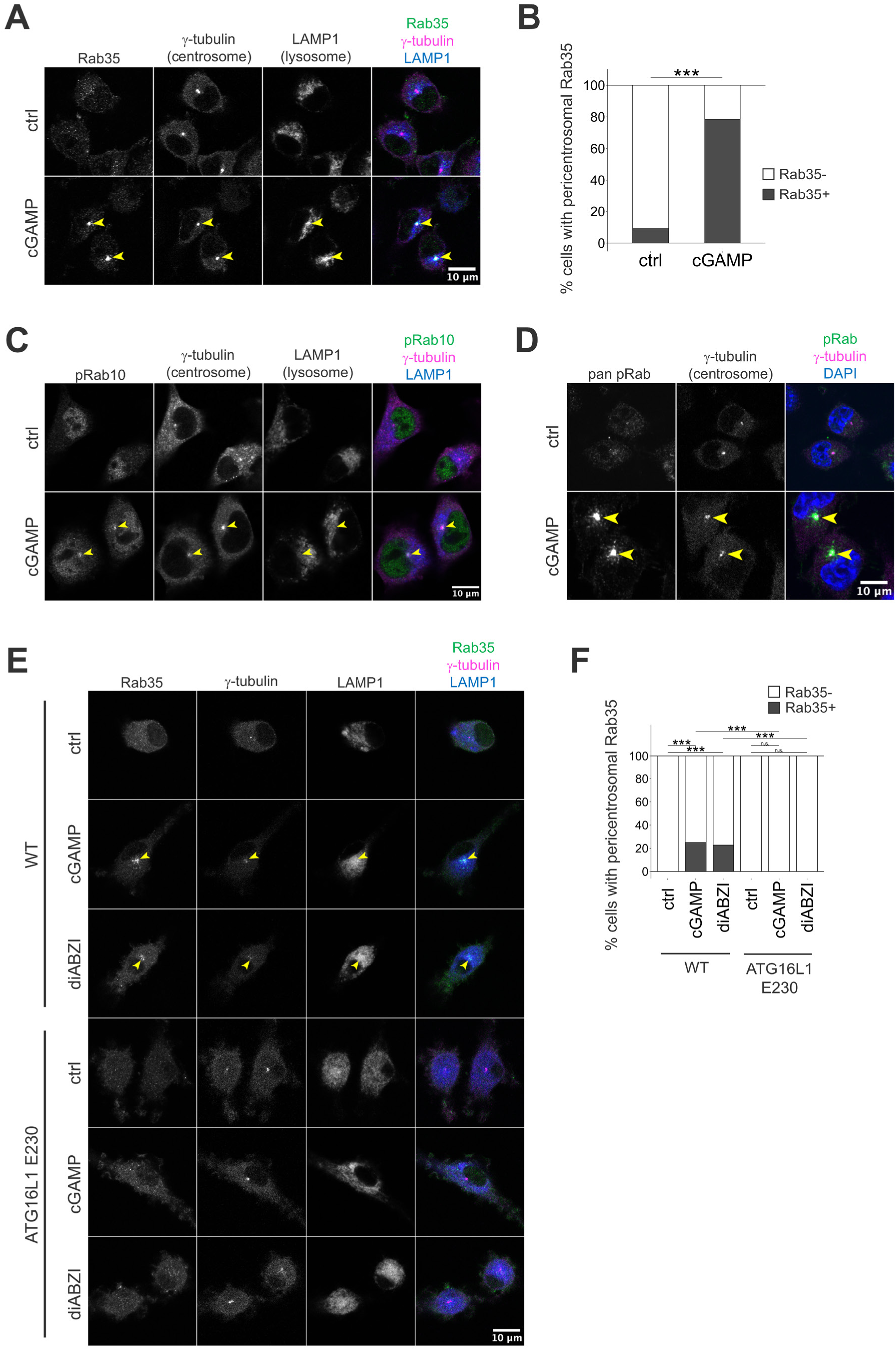
Rab proteins translocate to the pericentrosomal area upon STING activation in a VAIL-dependent manner. (**A**) Immunocytochemical analysis for Rab35 and indicated markers in MG6 cells treated with or without cGAMP. Arrowheads indicate Rab35-positive centrosomes. (**B**) Quantification of the percentages of cells harboring Rab35-positive centrosomes in A. Statistical significance was assessed by chi-squared tests with Holm-Bonferroni’s correction. (**C**, **D**) Immunocytochemical analysis for pRab10 (C), pan-pRab (D) in MG6 cells treated with or without cGAMP. Arrowheads indicate the centrosomes positive for each protein. (**E**) Immunocytochemical analysis for Rab35 and indicated markers in WT or ATG16L1 ΔWD40 BMDMs treated with cGAMP or diABZI. (**F**) Quantification of the percentages of cells harboring the Rab35-positive centrosome in E. Statistical significance was assessed by chi-squared tests with Holm-Bonferroni’s correction. n.s.: not significant, ***: *p* < 0.001.

We further examined whether Rab35 recruitment is regulated by VAIL using bone marrow-derived macrophages (BMDMs) prepared from wild-type (WT) or Atg16L1-ΔWD40 mice (E230 mice), in which a premature stop codon at Pro231 in Atg16l1 prevents translation of the C-terminal WD40 domain [21]. We found that WT BMDMs, but not ATG16L1-ΔWD40 BMDMs, exhibited Rab35 accumulation at the pericentrosomal area under treatment with cGAMP or diABZI (**Figures 6C,D**). Endogenous LRRK2 was below the detection limit by immunocytochemistry in BMDMs; we therefore showed Rab35 accumulation as a readout of the pathway in these cells. Together, these results indicate that Rab35 accumulates at the pericentrosomal area downstream of STING activation in a VAIL-dependent manner, and that Rab35 is required for the pericentrosomal translocation of LRRK2.

### Microtubule-based transport and motor adaptors are required for the pericentrosomal translocation of LRRK2

Since the centrosome is the major microtubule-organizing center in cells [73], we asked whether microtubule structure is required for the translocation of LRRK2. Treatment with the microtubule-depolymerizing agent nocodazole inhibited cGAMP-induced accumulation of phosphorylated Rab10 (**Figures 7A,B**) without affecting TBK1 phosphorylation (**Figures 7A,C**). In addition, the translocation of LRRK2 to the pericentrosomal area was suppressed by nocodazole treatment (**Figure 7D**).

**Figure 7.**
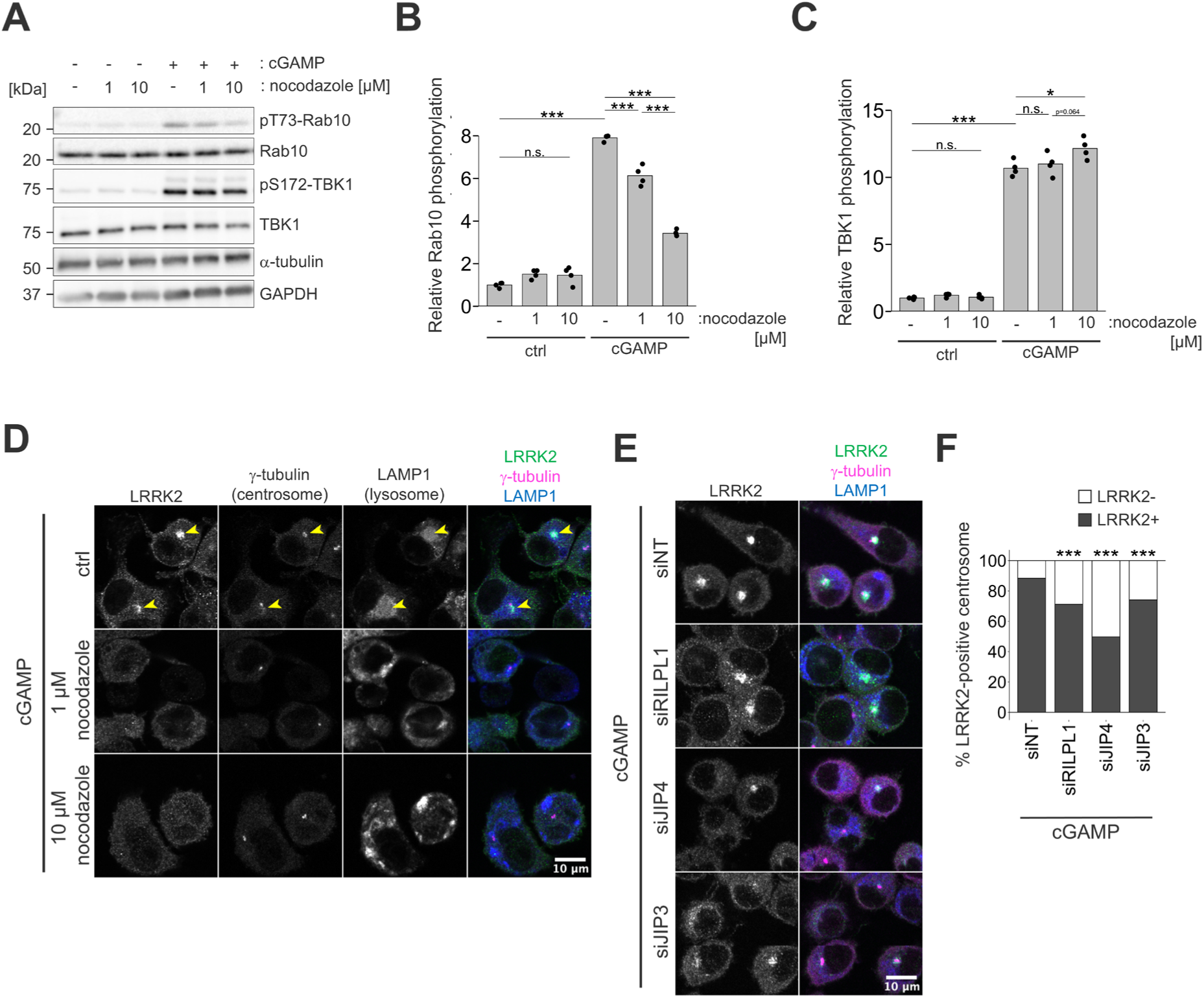
The pericentrosomal translocation of LRRK2 depends on microtubule-based transportation. (**A**-**C**) Immunoblotting of lysates of MG6 cells treated with nocodazole for two hours before cGAMP treatment. Rab10 phosphorylation and TBK1 phosphorylation were quantified in B and C, respectively. Statistical significance was assessed by two-way ANOVAs with Tukey’s post-hoc tests. (**D**) Immunocytochemical analysis of MG6 cells treated with nocodazole for two hours before cGAMP treatment. Arrowheads indicate the LRRK2-positive centrosomes. (**E**) Immunocytochemical analysis of RAW264.7 cells treated with siRNAs targeting the indicated adaptor proteins. (**F**) Quantification of the percentages of cells harboring the LRRK2-positive centrosome in E, scored as in Fig. 1G. Statistical significance was assessed by chi-squared tests with Holm-Bonferroni’s correction between siNT and each siRNA-treated sample. n.s.: not significant, *: *p* < 0.05, ***: *p* < 0.001.

Candidate mediators of microtubule-based transport downstream of the LRRK2-Rab35 axis include RILPL1, JIP4 and JIP3, which serve as motor adaptors and as effectors of LRRK2-phosphorylated Rab GTPases including Rab35 [39,41–43,53]. Knockdown of each adaptor significantly suppressed the translocation of LRRK2 to the pericentrosomal area, with JIP4 knockdown producing the strongest suppression (**Figures 7E,F**). Knockdown efficiencies were confirmed previously [37] or by RT-qPCR (**Fig. S4F,G**). This result accords with previous reports which suggested that Rab35 recruits JIP4 as its effector [39] or acts as a master Rab that concentrates multiple Rab proteins and their effectors, including JIP4, at recycling endosomes [74]. Yet, knockdown of RILPL1 or JIP3 also significantly suppressed the pericentrosomal translocation of LRRK2, suggesting redundant roles of these adaptor proteins. Together, these findings suggest that LRRK2 is recruited to V-ATPase-positive organelles upon STING activation, where LRRK2 promotes transport along microtubules via the motor adaptors.

We further examined whether stimuli other than STING could induce similar LRRK2-dependent changes in lysosomal positioning. We previously reported that LRRK2 is detected on enlarged lysosomes in RAW264.7 cells treated with chloroquine (CQ) [37], while only some of enlarged lysosomes were LRRK2-positive (**Fig. S5A**). We then hypothesized that LRRK2-positive lysosomes enlarged by CQ treatment are also accumulated toward the centrosome. We measured the distance from the centrosome to the nearest surface of each lysosome because they were enlarged and individually identifiable by CQ treatment. LRRK2-positive lysosomes were located significantly closer to the centrosome than LRRK2-negative ones within the same cells (**Fig. S5B,C**). These observations indicate that the association between LRRK2 and pericentrosomal positioning of lysosomes is not restricted to STING activation but also occurs under general stressors that induce VAIL.

### LRRK2 kinase activity is dispensable for the transcriptional responses downstream of STING

We examined whether LRRK2 activation affects well-studied STING-mediated immune response pathways. STING activation primarily upregulates transcription of *Ifnb1* (encoding interferon-β [IFNβ]) and interferon-stimulated genes (ISGs), which requires TBK1 and interferon regulatory factor 3 (IRF3) [75,76]. Treatment of RAW264.7 cells with cGAMP for six hours increased the expression of *Ifnb1*, whereas inhibition of LRRK2 kinase activity did not cause significant changes (**Fig. S6A**). The expression of *Isg15* and *Ifit1*, two representative ISGs, also increased upon cGAMP treatment, while MLi-2 treatment had no effect (**Fig. S6B,C**). The expression of *Ifna*, which is induced by IFNβ-mediated activation of IFNα/β receptor (IFNAR) [77], increased upon cGAMP treatment, while no effect of MLi-2 was observed (**Fig. S6D**). Thus, LRRK2 kinase activity does not appear to modify the typical type I interferon response induced by STING. STING also induces the production of pro-inflammatory cytokines independently of IRF3-based interferon signaling [78]. Treatment with cGAMP for six hours increased the expression of *Il6* and *Tnf*, while MLi-2 did not cause significant changes (**Fig. S6E,F**), suggesting that LRRK2 kinase activity does not modify STING-induced cytokine production. Together with the previous finding that TBK1 is dispensable for LRRK2 activation downstream of STING [44], these results suggest that the LRRK2 and TBK1 branches of STING signaling operate largely in parallel.

Recent studies have shown that STING activation promotes the nuclear translocation of MiT/TFE transcription factors (TFE3, TFEB, and MITF) and transcription of lysosomal genes [79,80]. This process has been shown to depend on VAIL and the lipidation of GABARAP family members [8,81], while mutational upregulation of LRRK2 kinase activity conversely has been shown to downregulate expressions of lysosomal genes [82]. We assessed the nuclear translocation of TFEB and TFE3 in MG6 cells under STING activation or mTORC1 inhibition (**Fig. S7A,B**). Nuclear TFEB and TFE3 signals were significantly enriched after treatment with cGAMP, while inhibition of LRRK2 kinase activity by MLi-2 did not affect their nuclear translocation (**Fig. S7C,D**). Although cGAMP induced nuclear translocation of TFEB and TFE3, we did not detect a corresponding induction of lysosomal gene transcripts other than a modest increase in *Lamp1* (**Fig. S7E–H**). This differs from previous reports in other cell types [79,80] and may reflect cell type-specific differences or the time points examined.

Thus, LRRK2 kinase activity is dispensable for the transcriptional arm of STING signaling, including type I interferon responses, pro-inflammatory cytokine induction, and MiT/TFE-dependent lysosomal gene expression, at least under the conditions examined here.

### Enhanced LRRK2 kinase activity promotes pericentrosomal accumulation of GABARAP and Rab35 upon STING activation

Since mutations in LRRK2 enhance kinase activity and cause familial PD, we sought to examine whether the G2019S mutation affects STING-responsive VAIL, LRRK2, and the translocation of related proteins to the pericentrosomal area. Primary microglia were prepared from WT or LRRK2-G2019S bacterial artificial chromosome (BAC) transgenic (Tg) mice and treated with cGAMP. We first confirmed the enhanced phosphorylation of TBK1 and dimerization of STING by cGAMP treatment in both WT and G2019S-Tg cells (**Figures 8A,B**). In G2019S-Tg cells, Rab10 phosphorylation was elevated under untreated conditions and was further enhanced by cGAMP treatment (**Figure 8A**). LRRK2 was detected at the pericentrosomal area in cGAMP-treated G2019S-Tg microglia, confirming that this translocation also occurs in primary microglia carrying a pathogenic LRRK2 mutation (**Figures 8C,F**). Because BAC transgenic microglia express LRRK2 well above endogenous levels (**Figure 8A**), the difference in LRRK2 signal between genotypes cannot be attributed to the mutation itself. We therefore used Rab35 and GABARAP, which are expressed at endogenous levels in both genotypes (**Figure 8A**), as readouts of the pathway. Pericentrosomal Rab35 was detected in a subset of WT microglia under basal conditions, and its frequency increased after cGAMP treatment (**Figures 8D,G**). G2019S-Tg cells showed a higher frequency than WT cells both before and after stimulation. Pericentrosomal GABARAP was detectable only in cGAMP-treated G2019S-Tg cells and was below the detection threshold in the other three conditions (**Figures. 8E,H**).

**Figure 8.**
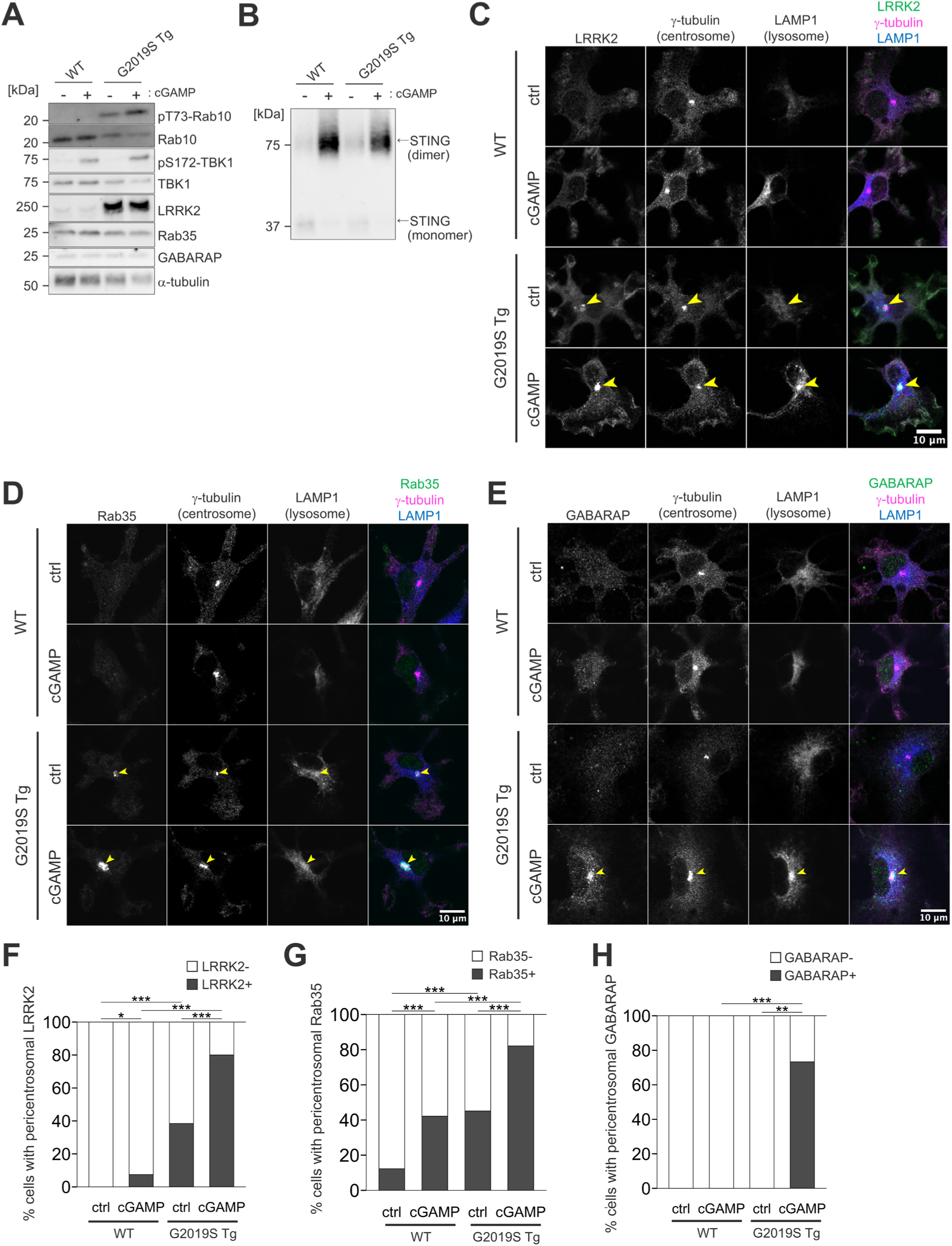
Hyperactivation of LRRK2 induces upregulation of the events directing organelle to the pericentrosomal area. (**A**) Immunoblotting of lysates of primary microglia prepared from WT or *Lrrk2* G2019S Tg pups, primed by IFNγ and treated with cGAMP for six hours. (**B**) Non-reducing immunoblotting of the same sets of cell lysates as in A probed with an anti-STING antibody. (**C**-**E**) Immunocytochemical analysis of primary microglia treated with or without cGAMP. Arrowheads indicate the centrosomes positive for LRRK2, Rab35, and GABARAP. (**F**-**H**) Quantification of the percentages of cells harboring the centrosome positive for LRRK2, Rab35, or GABARAP. Statistical significance was assessed by chi-squared tests with Holm-Bonferroni’s corrections. *: *p* < 0.05, **: *p* < 0.01, ***: *p* < 0.001.

These results suggest that, under conditions where LRRK2 kinase activity is enhanced, the accumulation of VAIL-induced proteins at the pericentrosomal area is facilitated. Interestingly in G2019S-Tg cells, pericentrosomal Rab35 was detected in a substantial fraction of cells even without cGAMP stimulation (**Figures 8D,G**), whereas this was rare in WT cells. This suggests that enhanced LRRK2 kinase activity lowers the threshold for engaging the LRRK2–Rab35 axis, such that the constitutive level of VAIL becomes sufficient to drive pericentrosomal translocation without additional STING-mediated stimulation.

Overall, our findings define a pathway in which STING activation induces VAIL on lysosomes and REs, recruits LRRK2 to these organelles, and drives their microtubule-based transport toward the centrosome through Rab35 and its motor adaptors (**Figure 9**). The G2019S mutation, which enhances LRRK2 kinase activity, facilitates this pathway.

**Figure 9.**
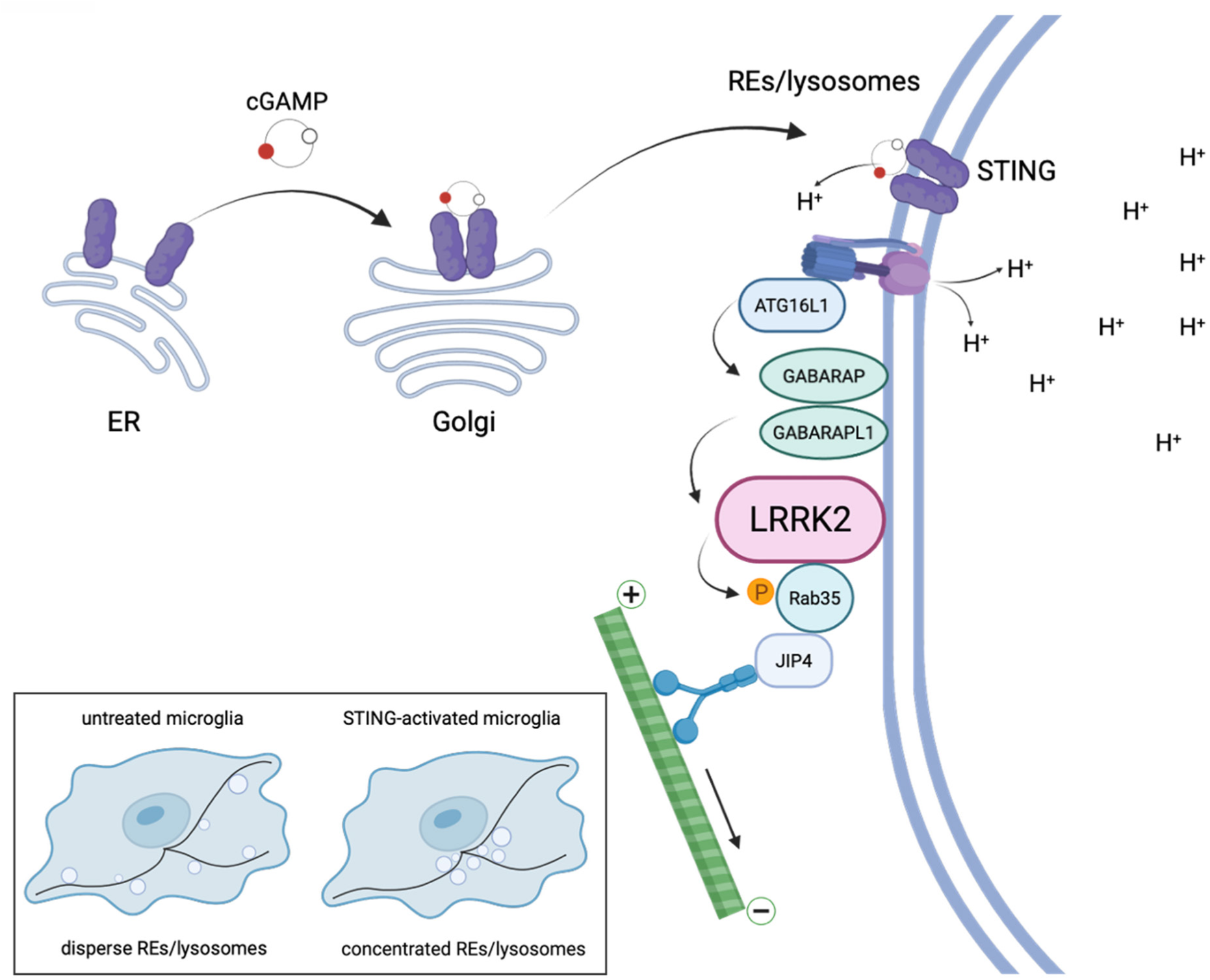
Schematic depiction of pericentrosomal positioning of acidic organelles via the STING-VAIL-LRRK2-Rab35 axis. A cartoon depicting the STING-VAIL-LRRK2-Rab35 axis to modify lysosomal subcellular distribution. STING activation induces VAIL at lysosomes and REs, then LRRK2 is recruited to these organelles. LRRK2 employs Rab35 and adaptor proteins including JIP4 by its kinase activity, facilitating positioning of LRRK2-positive lysosomes and REs toward the pericentrosomal area.

## Discussion

In this study, we found that STING activation induces LRRK2 activation and steers the acidic organelles to concentrate at the pericentrosomal area in a LRRK2-dependent manner. This phenomenon was observed as the formation of unique puncta at the pericentrosomal area that contain LRRK2, VAIL-related proteins (LC3B, GABARAP and ATG16L1), Rab GTPases and acidic organelles. This pathway appears to be driven independently of known STING downstream mechanisms such as innate immune responses and lysosomal biogenesis, suggesting that STING activation regulates diverse cellular functions by simultaneously activating multiple signaling pathways.

LRRK2 itself has only rarely been observed at the centrosome. Earlier work using a BAC-based LRRK2 reporter noted that a fraction of LRRK2-positive puncta lay close to the γ-tubulin-positive centrosome, in a study focused on the autophagy-lysosome system [30]. Subsequent studies detected pericentrosomal or filamentous microtubule-related LRRK2 only under specific conditions: overexpression of pathogenic variants such as G2019S, R1441C or Y1699C, or pharmacological kinase inhibition, both of which relocalize LRRK2 to the centrosome or to microtubule-associated filamentous structures [55,83,84]. Wild-type LRRK2 remained largely cytosolic in these settings, and the associated centrosomal phenotypes were attributed to phosphorylated Rab8A and Rab10 rather than to LRRK2 itself [53,55]. Our findings differ from these reports in three aspects: first, the pericentrosomal accumulation we describe involves endogenous, wild-type LRRK2; second, it is triggered by physiological stimuli rather than by mutation or overexpression; third, it requires, rather than follows, LRRK2 kinase activity. We therefore consider the STING-induced translocation of LRRK2 to be mechanistically distinct from the centrosomal association observed under overexpression or kinase inhibition, and its detection in microglia and macrophages may reflect a property of these cell types that is not shared by the cell lines used in earlier work.

Upon STING activation, the pericentrosomal area was positive for lysosomes and REs, both of which are involved in STING signal termination [64–66]. Consistent with this, LRRK2 observed at the pericentrosomal area was detected in fractions containing lysosomes and REs. However, we have not been able to clearly identify which organelle LRRK2 is recruited to, as the lysosomal and RE fractions partially overlap. A previous study localized LRRK2 to lysosomal fractions purified with superparamagnetic iron oxide nanoparticles [44], although such preparations may also contain endosomal compartments. Since STING has been reported to reside on REs during the termination of STING signaling [64], it is possible that REs are the site of VAIL and the destination of LRRK2 recruitment as well as lysosomes.

STING has been reported to function as a conserved proton channel, and its activation results in mild deacidification of the Golgi apparatus and post-Golgi trafficking vesicles [16,17,85]. Thus, the transport of LRRK2-positive lysosomes and REs toward the centrosome may function as a response to deacidifying stress. Notably, lysosomes are known to exhibit slight differences in acidity depending on their intracellular location, with those at the cell periphery being less acidic and having lower protease activity than those near the centrosome [86]. Given that VAIL depends on the activity of the proton pump V-ATPase, this lysosome/RE transport toward the centrosome appears to be a reasonable response to selectively act on stressed organelles under deacidifying stress. This idea is also supported by a report observing JIP4-dependent lysosomal retrograde transport in response to oxidative stress [87].

LRRK2 kinase activity could potentially function both as a cause and a consequence of the pericentrosomal translocation of LRRK2/lysosomes. Endolysosomal compartments with substrate Rabs on their surface are mainly enriched in the pericentrosomal region [88,89], whereas LRRK2-phosphorylated Rab35 on lysosomes is thought to direct themselves toward the centrosome, thereby enabling the interaction between LRRK2 and the Rab pool. This mechanism may explain the enrichment of phosphorylated Rabs at the pericentrosomal region. This model is supported by the prior finding that peripheral targeting of lysosomes decreases Rab10 phosphorylation, whereas perinuclear targeting enhances it [90]. Additionally, decrease in Rab10 phosphorylation under nocodazole treatment was observed not only in this study but also in a recent study [91]. Although the latter study focused more on the role of cellular GTP, these studies support the notion that microtubules are required for LRRK2 to reach its substrate pool.

We also found that under CQ treatment, LRRK2-positive enlarged lysosomes were significantly closer to the centrosome compared with LRRK2-negative ones. This observation can be explained either by the selective movement of LRRK2-positive lysosomes occurring similarly to STING activation, or by the greater likelihood of LRRK2 recruitment via VAIL occurring near the centrosome. Overall, this suggests that the mechanism controlling organelle positioning through the VAIL-LRRK2 pathway operates beyond the context of STING activation.

The possibility that the STING-LRRK2 pathway functions beyond organelle positioning also remains. Pericentrosomal Rab proteins phosphorylated by LRRK2 have been shown to cause the impairment in primary ciliogenesis in various cell types or centrosomal cohesion deficits in dividing cells [23,53,55,92]. Although these phenotypes were not observed in our microglia even under STING activation, they may be observed in other cell types under such conditions. On the other hand, pericentrosomal localization of LRRK2 and acidic organelles that we discovered here may be easily detected in microglia and macrophage-lineage cells. Therefore, it might be reasonable to speculate that the STING-LRRK2 pathway plays distinct roles depending on the cell type.

Regarding the involvement of VAIL in the recruitment of LRRK2 upon STING activation, GABARAP and GABARAPL1 were found to strongly contribute to the process, concurring the previous studies that have shown GABARAP to be critical for VAIL-mediated LRRK2 activation [8,44]. Several Rab proteins have been reported to modify microtubular transportation via kinesin/dynein adaptor proteins [39,41,42,53]; for Rab35, JIP4 was previously described as a downstream effector resident on lysosomes [39] or recycling endosomes [74]. Thus, our findings that the series of events are primarily driven by Rab35-JIP4 axis appear to be valid considering previous reports, although RILPL1 and JIP3 were suggested to have redundant roles.

Several limitations lie in this study. First, the spatial resolution of confocal microscopy was not sufficient to resolve LRRK2-positive organelles from the centrosome itself; super-resolution or electron microscopy will be required to fully define this relationship. Second, because RAW264.7 and MG6 cells are incompetent to transfection-based overexpression, rescue experiments for the knockdown phenotypes were not feasible. Third, BafA1 treatment inhibits the V-ATPase globally and thereby affects lysosome positioning and mTORC1 signaling independently of VAIL, preventing it from further solidifying the role of VAIL in lysosome positioning. Finally, the LRRK2-G2019S BAC transgenic microglia express LRRK2 above endogenous levels, so genotype comparisons were restricted to downstream components whose expression was unaffected.

In summary, our findings suggest STING’s unconventional role in the spatial reorganization of lysosomes and REs in microglial cells by employing LRRK2 via VAIL. This molecular cascade is suggested to arise not only in response to deacidifying stress associated with STING activation, but also in response to other VAIL-inducing stresses such as CQ treatment. The physiological purpose of this repositioning remains to be defined. It will be important to determine the consequences of aberrant activation of this pathway in vivo, especially its contribution to the pathogenesis of LRRK2-associated PD and other diseases involving the cGAS-STING pathway, using appropriate disease models and other approaches.

## Methods

### Cell culture and drug treatments

MG6 cells and RAW264.7 cells were cultured in a 10 cm dish for suspended cells (Sumitomo Bakelite Co., Ltd., MS-1390R) in DMEM (FUJIFILM Wako Pure Chemical Corporation, 044-29765) supplemented with 10% (v/v) FBS (Biowest, S1400-500) and 1% (v/v) penicillin/streptomycin (FUJIFILM Wako, 168-23191). Before experiments, cells were seeded on 24-well plates (AGC Techno Glass Co., Ltd., 3820-024) and always primed by mouse IFNγ (15 ng/mL, Cell Signaling Technology, 39127S) unless otherwise noted. For STING activation, 2′,3′-cGAMP (final 10 µg/mL, Selleck Chemicals, S7904) or diABZI (final 1 µM, Selleck Chemicals, S8796) was added to the medium for six hours unless otherwise mentioned. For VAIL activation, SaliP (final 500 nM, Omm Scientific, Inc.) was added to the medium for three hours. MLi-2 (final 100 nM, Abcam, ab254528), H-151 (final 5 µM, Selleck Chemicals, S6652) and bafilomycin A_1_ (final 100 nM, MedChemExpress, HY-100558) were used for the inhibition of LRRK2, STING and V-ATPase, respectively. For microtubule destabilization, nocodazole (final 1 or 10 µM, Sigma-Aldrich, M1404) was added two hours before cGAMP treatment and was left in the culture media throughout the treatment.

### Knockdown treatment

For knockdown experiments, unprimed RAW264.7 cells seeded on 24-well plates were transfected with final 15 nM of each siRNA (Dharmacon siGENOME, Revvity) using Lipofectamine™ RNAiMAX (Thermo Fisher Scientific, 13778-150) according to the manufacturer’s protocol. Catalog id for each siRNA is shown in **Table S1**. Cells were then cultured for 72 hr, with IFNγ added in the last 24 hr. For knockdown of multiple genes, multiple siRNAs were mixed and transfected so that the total amount of siRNA remained the same. Knockdown efficiencies were confirmed previously or by RT-qPCR analysis, as shown in **Fig. S4**.

### Antibodies

The following antibodies were used: anti-LRRK2 (Abcam, MJFF2 [c41-2]), anti-LRRK2 (Epitomics, [UDD 3 30 (12)]), anti-phospho-Thr73 Rab10 (Abcam, MJF-R21 [ab230261]), anti-Rab10 (Cell Signaling Technology, D36C4), anti-phospho-Rab (Abcam, anti-phospho-Thr72 Rab8A [MJF-R20]), anti-α-tubulin (Sigma-Aldrich, DM1A), anti-γ-tubulin (Abcam, GTU-88 [ab11316]), anti-mouse LAMP1 (Bio-Rad Laboratories, 1D4B), anti-Rab11 (Cell Signaling Technology, D4F5), anti-TGN46 (Abcam, ab16059), anti-EEA1 (Cell Signaling Technology, 2411), anti-LC3B (Sigma-Aldrich, L7543), anti-ATG16L1 (Cell Signaling Technology, D6D5), anti-GABARAP (Cell Signaling Technology, E1J4E), anti-Rab35 (Proteintech Group, 11329-2-AP), anti-TFEB (Proteintech Group, 13372-1-AP), anti-TFE3 (Cell Signaling Technology, #81744), and anti-ARL13B (NeuroMab Facility, 75-287-020).

### Immunocytochemistry

Cells were seeded on coverslips (Matsunami Glass Ind., Ltd., C012001) in a 24-well plate and primed with IFNγ for >18 hours, then treated with each drug. Cells were fixed with prewarmed 4% paraformaldehyde (TAAB Laboratories Equipment Ltd., p001) in PBS for 30 minutes followed by washing with prewarmed DPBS. Cells were blocked and permeabilized by blocking solution (3% BSA [Sigma-Aldrich or FUJIFILM Wako, A2153-100G or 013-27054] and 0.1% Triton X-100 [MP Biomedicals, LLC, 194854] in DPBS) for an hour. Cells were incubated with primary antibodies (2 hr, RT, used in 500× dilution for LAMP1, 100× for γ-tubulin, and 200× for others) followed by one hour incubation by Alexa-conjugated secondary antibodies (Invitrogen, A-11034, A-21206, A-11030, and A-21247, used in 500× dilution) and DAPI (Invitrogen, D1306) both in blocking solution. After staining, cells were mounted with hydrophilic mountant (Beckman Coulter, Inc., PN IM0752).

### Confocal microscopy and image preparation

Immunostained samples were observed by confocal microscope (Evident, FV3000) using 100× objective lens. For each condition, three or more regions composed of 5×5 tiles (611 µm square after stitching) were randomly chosen to be photographed. Boxed regions in Fig. 2A, 2B, 2G and 2H are magnified four times and shown using bicubic interpolation (Fiji/ImageJ 2.14.0/v1.54f).

### Quantification of images

Each stitched image was analyzed by a program written in Julia to calculate the frequency of accumulation of each protein at the pericentrosomal area and lysosome-centrosome proximity distance. All quantification was performed on the original, unscaled images. Briefly, nuclear and cytoplasmic regions were first determined, and then each cytoplasmic region was assigned to each nucleus. The use of the watershed method made this possible even in areas where cells have contact with each other. The centrosome region was then determined from γ-tubulin-stained images. Cells with the centrosome observed in the cytoplasmic region and not obscured by the edges of the image were selected for measurement. Most MG6 or RAW264.7 cells had centrosomes marked with γ-tubulin, but cells with centrosomes out of the focal plane or with centrosomes overlapping within the nuclear staining-positive area were excluded.

Pericentrosomal accumulation of LRRK2, LC3B, and Rab35 was scored per cell by comparing the mean intensity within 1 µm of the centrosome with that of the remaining cytoplasm. Cells with a ratio of ≥2 were scored as positive.

For the measurement of intracellular distribution of lysosomes or REs, we introduced the index of organelle-centrosome average distance, which is a weighted average of the distance from the centrosome to the pixels of LAMP1/Rab11 staining, as shown in the following equation (L: marker intensity at the pixel, d: distance between the pixel and the centrosome).

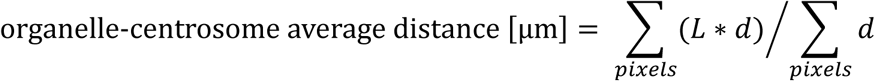

One-way ANOVA or two-way ANOVA with Tukey’s post-hoc test was conducted to verify the difference in organelle-centrosome proximity distance among conditions.

For the quantification of the distance between the centrosome and the enlarged lysosomes after CQ treatment, the shortest distance from each lysosome’s edge to the centrosome was measured as distance, as shown in Fig. S5B.

For quantification of nuclear enrichment of TFEB or TFE3, the nuclear signal was defined as the 80th percentile (p80) intensity within the nucleus, eroded by 2 px to exclude the boundary, and the cytoplasmic signal as the p80 intensity within a perinuclear ring located 2–4 px outside the nuclear boundary (i.e. a 2 px gap was left at the boundary to avoid spillover). The nuclear enrichment index was calculated as the ratio of nuclear to ring p80.

### Immunoblotting

Cells were seeded and primed with IFNγ for >18 hours, then treated with each drug. Cells were collected by ice-cold lysis buffer (1% Triton X-100 in Tris-buffered saline (TBS) supplemented with cOmplete™ [Roche Diagnostics GmbH, 05056489001, one tablet per 50 mL], EDTA-free protease inhibitor cocktail and PhosSTOP™ [Roche Diagnostics GmbH, 4906837001, one tablet per 10 mL], phosphatase inhibitor cocktail) after washing with prewarmed DPBS. Cell debris was removed from the lysate by centrifugation (12,600 × *g*, 5 min), and the supernatants were mixed with NuPAGE™ LDS sample buffer (4×) (Thermo Fisher Scientific, NP0008) supplemented with 4% β-mercaptoethanol (FUJIFILM Wako, 139-07525) at a ratio of 3:1 (v/v) followed by incubation for five minutes at 95°C. When assessing STING dimerization, β-mercaptoethanol was not added to maintain non-reducing conditions, as shown previously [50]. Samples were then subjected to SDS-PAGE using polyacrylamide gels (SuperSep Ace, FUJIFILM Wako, 194-15021 [gradient], 190-15001 [fixed-percentage]) and transferred onto PVDF membranes (Invitrogen, IB34001) using iBlot3 (Invitrogen). Transferred membranes were blocked with 5% skim milk (BD Biosciences, 232100) + 0.1% Tween 20 in TBS for 30 minutes at RT. Membranes were incubated with primary antibodies (o/n, 4°C, used in 500× dilution for Rab10, 1,000× for others) in Immuno-enhancer Reagent A (FUJIFILM Wako, 091-05811) followed by incubation with HRP-conjugated secondary antibodies (Jackson ImmunoResearch Laboratories, Inc., 111-035-003 and 515-035-003, 45 min, RT, used in 5,000× dilution) in Immuno-enhancer Reagent B (FUJIFILM Wako, 098-05821). Protein bands were visualized by chemiluminescence using ImmunoStar® Zeta (FUJIFILM Wako, 291-72401) and detected using ImageQuant™ LAS-4000 (FUJIFILM Wako). For densitometric analysis, the integrated densities of protein bands were calculated using Fiji/ImageJ 2.14.0/v1.54f. To further align the overall chemiluminescence intensity among several different membranes, the band intensities were normalized so that the sum of the intensities of all bands for each membrane was equal.

### Sucrose density gradient centrifugation

MG6 cells were seeded on 6-well plates (AGC Techno Glass Co., Ltd., 3810-006N) and primed with IFNγ for >18 hours, then treated with or without cGAMP for six hours. Cells were washed by ice-cold DPBS and collected by a scraper. Samples were homogenized on ice by pumping by 1 mL syringe (Terumo Corporation, SS-01T) with 27-gauge needles (Terumo Corporation, NN-2719S) for 15 strokes. A buffer containing 0.25-2 M sucrose (FUJIFILM Wako, 196-00015) was layered in an ultracentrifuge tube, and the sample was placed on top. The mixture was then ultracentrifuged at 210,000 × *g* for three hours at 4°C with a swing rotor (Beckman Coulter, Inc., SW 41 Ti). Subsequently, 750 µL of aliquots were carefully collected from the low-density side as separated fractions and analyzed by immunoblotting.

### RT-qPCR

Cells were seeded on 24-well plates, primed with IFNγ for >18 hours, and treated with the indicated reagents. RNA was extracted using PureLink® RNA Mini Kit (Thermo Fisher Scientific, 12183018A), and reverse-transcribed with oligo(dT) primers using SuperScript™ III First-Strand Synthesis System (Invitrogen, 18080051) or ReverTra Ace® qPCR RT Master Mix with gDNA Remover (TOYOBO Co., Ltd., FSQ-301). Transcript levels of selected targets and a reference gene (*Gapdh*) were quantified by qPCR using LightCycler 480 system (Roche Diagnostics GmbH) or QuantStudio® 3 (Thermo Fisher Scientific). Primer sequences are listed in **Table S2**. Briefly, reactions were performed in a final volume of 20 µL containing 10 µL of LightCycler 480 SYBR Green I Master Mix (2×) (Roche Diagnostics GmbH, 04707516001), 0.08 µL (50 µM) of each primer, 1 µL (∼1 µg) cDNA, and 8.84 µL of water. The temperature profiles comprised an initial denaturation step at 95°C for 5 min followed by 45 cycles consisting of denaturation at 95°C for 10 sec, annealing at 60°C for 10 sec, and extension and data acquisition at 72°C for 10 sec. Amplified products were subjected to melting curve analyses. Relative expression levels normalized for *Gapdh* were calculated by the 2^-ΔΔCt method assuming 100% PCR efficiency (E = 2 per cycle).

### Primary microglia preparation

The isolation of primary microglia from mice was performed in accordance with the regulations and guidelines of the University of Tokyo and approved by the institutional review committee. LRRK2 G2019S Bac Tg mouse strain with C57BL/6J background (The Jackson Laboratory, #012467) were used for microglia isolation. The early postnatal (P0-P3) littermates were prepared by crossbreeding a wild-type C57BL/6J female with a G2019S TG heterozygous male. Primary microglia were isolated from each pup as reported [93,94]. Briefly, the cerebrum, with the meninges removed, was collected from each pup in ice-cold HBSS (FUJIFILM Wako, 085-09355) and triturated by repeated incubation with trypsin and pipetting. Tail of each pup was cut for genotyping. Cells were added to DMEM supplemented with 10% (vol/vol) FBS and 1% penicillin/streptomycin (DMEM (+/+)) and passed through a 100 µm strainer (AS ONE Corporation, VCS-100). Cells were spun, resuspended in DMEM (+/+), and seeded in T-75 flask (Thermo Fisher Scientific, one flask per pup). The culture medium was refreshed every three days. Fourteen days after seeding, microglia were isolated by shaking and tapping the flask to bring them to the surface, and were centrifuged at 2,000 rpm for five minutes. Cells were resuspended in medium supplemented with 10% L929 cell culture supernatant and cultured on plates or coverslips for two days until use.

### Statistical analysis

All data are expressed as mean ± s.e.m. Statistical details for each experiment, including the test used, the number of replicates and the definition of the statistical unit, are provided in the corresponding figure legends. Pairwise comparisons between two conditions, including the quantification of immunoblots, the LCAD comparison, the confirmation of knockdown efficiencies, and the quantification of nuclear translocation (ICC), were performed using unpaired two-tailed Student’s *t*-tests. Comparisons of STING dimerization between conditions were performed using unpaired Welch’s *t*-tests. Experiments comprising a single experimental factor with more than two levels, such as quantification of immunoblots, were analyzed by one-way ANOVA followed by Tukey’s post-hoc test. Experiments comprising two experimental factors, including quantifications of immunoblots, of LCADs, and of RT-qPCR data, were analyzed by two-way ANOVA followed by Tukey’s post-hoc test. Frequency data, namely the proportion of cells displaying centrosomal localization of the indicated proteins, were analyzed by chi-squared tests.

Where multiple *t*-tests or chi-squared tests were applied to the same dataset, *p* values were adjusted using the Holm-Bonferroni method. A *p* value below 0.05 was considered statistically significant. \*\*\**p* < 0.001, \*\**p* < 0.01, \**p* < 0.05. Analyses were performed using R 4.3.1, Julia 1.10.0, or Microsoft Excel 16.112.1.

## Data availability

All data sets used or analyzed in this study are available from the corresponding author upon request.

## Acknowledgements

We thank Drs. Thomas Wileman and Matthew Jefferson (University of East Anglia, United Kingdom) for providing bone marrow-derived macrophages from wild-type and *Atg16l1*-ΔWD40 mice, Megumi Kurosu (The American School in Japan) for assistance with image analysis, and members of the Iwatsubo laboratory for helpful suggestions and discussions. Images in Figure 3A and Figure 9 were created in BioRender. Suenaga, S. (2026) https://BioRender.com/s6wk2do. This study was supported by JSPS KAKENHI grant numbers 22H02949 (T. K.), 23K24210 (T. K.), 25K02458 (T. K.), 24KJ0870 (S. S.), 23K27515 (T. I.), Takeda Science Foundation (T. K.), and ONO Medical Research foundation (T. K.).

## Author contributions

Conceptualization: S. S. and T. K.; Investigation: S. S., T. K. and M. S.; Methodology: S. S., T. K. and M. S.; Data curation: S. S., T. K. and M. S.; Writing – original draft: S. S. and T. K.; Writing – review & editing: T. K., M. S. and T. I.; Supervision: T. K.; Project administration: T. K.; Funding acquisition: S. S., T. K. and T. I. All authors read and approved of the final manuscript.

## Conflict of interest

The authors declare no competing financial interests.

## Notes

### Competing Interest Statement

The authors have declared no competing interest.

## References

[1] S. Dvorkin, S. Cambier, H.E. Volkman and D.B. Stetson, New frontiers in the cGAS-STING intracellular DNA-sensing pathway, Immunity 57 (2024), pp. 718–730.

[2] Y. Xiao, Y. Ma, J. Zhao, X. Zhang, C. Gan, J. Gao et al., The cGAS-STING signaling pathway as a therapeutic target in human diseases, Chin. Med. J. (Engl.) 138 (2025), pp. 3251–3284.

[3] T. Abe, A. Harashima, T. Xia, H. Konno, K. Konno, A. Morales et al., STING recognition of cytoplasmic DNA instigates cellular defense, Mol. Cell 50 (2013), pp. 5–15.

[4] Q. Chen, L. Sun and Z.J. Chen, Regulation and function of the cGAS-STING pathway of cytosolic DNA sensing, Nat. Immunol. 17 (2016), pp. 1142–1149.

[5] K.V. Swanson, R.D. Junkins, C.J. Kurkjian, E. Holley-Guthrie, A.A. Pendse, R. El Morabiti et al., A noncanonical function of cGAMP in inflammasome priming and activation, J. Exp. Med. 214 (2017), pp. 3611–3626.

[6] K.R. Balka, C. Louis, T.L. Saunders, A.M. Smith, D.J. Calleja, D.B. D’Silva et al., TBK1 and IKKε act redundantly to mediate STING-induced NF-κB responses in myeloid cells, Cell Rep. 31 (2020), pp. 107492.

[7] T.D. Fischer, C. Wang, B.S. Padman, M. Lazarou and R.J. Youle, STING induces LC3B lipidation onto single-membrane vesicles via the V-ATPase and ATG16L1-WD40 domain, J. Cell Biol. 219 (2020),.

[8] T. Huang, C. Sun, F. Du and Z.J. Chen, STING-induced noncanonical autophagy regulates endolysosomal homeostasis, Proc. Natl. Acad. Sci. U. S. A. 122 (2025), pp. e2415422122.

[9] X. Gui, H. Yang, T. Li, X. Tan, P. Shi, M. Li et al., Autophagy induction via STING trafficking is a primordial function of the cGAS pathway, Nature 567 (2019), pp. 262–266.

[10] K.M. Hooper, E. Jacquin, T. Li, J.M. Goodwin, J.H. Brumell, J. Durgan et al., V-ATPase is a universal regulator of LC3-associated phagocytosis and non-canonical autophagy, J. Cell Biol. 221 (2022),.

[11] J. Durgan and O. Florey, Many roads lead to CASM: Diverse stimuli of noncanonical autophagy share a unifying molecular mechanism, Sci. Adv. 8 (2022), pp. eabo1274.

[12] K.B. Boyle, C.J. Ellison, P.R. Elliott, M. Schuschnig, K. Grimes, M.S. Dionne et al., TECPR1 conjugates LC3 to damaged endomembranes upon detection of sphingomyelin exposure, EMBO J. 42 (2023), pp. e113012.

[13] D.P. Corkery, S. Castro-Gonzalez, A. Knyazeva, L.K. Herzog and Y.-W. Wu, An ATG12-ATG5-TECPR1 E3-like complex regulates unconventional LC3 lipidation at damaged lysosomes, EMBO Rep. 24 (2023), pp. e56841.

[14] N. Kaur, L.R. de la Ballina, H.S. Haukaas, M.L. Torgersen, M. Radulovic, M.J. Munson et al., TECPR1 is activated by damage-induced sphingomyelin exposure to mediate noncanonical autophagy, EMBO J. 42 (2023), pp. e113105.

[15] Y. Wang, M. Jefferson, M. Ramos, M. Whelband, K. Kreuzer, G. Khuu et al., The TECPR1:ATG5-ATG12 complex conjugates LC3/ATG8 to damaged lysosomes that expose luminal glycans in response to osmotic imbalance, Autophagy Rep. 4 (2025), pp. 2476218.

[16] B. Liu, R.J. Carlson, I.S. Pires, M. Gentili, E. Feng, Q. Hellier et al., Human STING is a proton channel, Science 381 (2023), pp. 508–514.

[17] J. Xun, Z. Zhang, B. Lv, D. Lu, H. Yang, G. Shang et al., A conserved ion channel function of STING mediates noncanonical autophagy and cell death, EMBO Rep. 25 (2024), pp. 544–569.

[18] J.-Y. Kim, H. Zhao, J. Martinez, T.A. Doggett, A.V. Kolesnikov, P.H. Tang et al., Noncanonical autophagy promotes the visual cycle, Cell 154 (2013), pp. 365–376.

[19] O. Florey, N. Gammoh, S.E. Kim, X. Jiang and M. Overholtzer, V-ATPase and osmotic imbalances activate endolysosomal LC3 lipidation, Autophagy 11 (2015), pp. 88–99.

[20] K. Fletcher, R. Ulferts, E. Jacquin, T. Veith, N. Gammoh, J.M. Arasteh et al., The WD40 domain of ATG16L1 is required for its non-canonical role in lipidation of LC3 at single membranes, EMBO J. 37 (2018), pp. e97840.

[21] S. Rai, M. Arasteh, M. Jefferson, T. Pearson, Y. Wang, W. Zhang et al., The ATG5-binding and coiled coil domains of ATG16L1 maintain autophagy and tissue homeostasis in mice independently of the WD domain required for LC3-associated phagocytosis, Autophagy 15 (2019), pp. 599–612.

[22] M. Steger, F. Tonelli, G. Ito, P. Davies, M. Trost, M. Vetter et al., Phosphoproteomics reveals that Parkinson’s disease kinase LRRK2 regulates a subset of Rab GTPases, Elife 5 (2016),.

[23] M. Steger, F. Diez, H.S. Dhekne, P. Lis, R.S. Nirujogi, O. Karayel et al., Systematic proteomic analysis of LRRK2-mediated Rab GTPase phosphorylation establishes a connection to ciliogenesis, Elife 6 (2017),.

[24] C. Paisán-Ruíz, S. Jain, E.W. Evans, W.P. Gilks, J. Simón, M. van der Brug et al., Cloning of the gene containing mutations that cause PARK8-linked Parkinson’s disease, Neuron 44 (2004), pp. 595–600.

[25] A. Zimprich, S. Biskup, P. Leitner, P. Lichtner, M. Farrer, S. Lincoln et al., Mutations in LRRK2 cause autosomal-dominant parkinsonism with pleomorphic pathology, Neuron 44 (2004), pp. 601–607.

[26] A.B. West, D.J. Moore, S. Biskup, A. Bugayenko, W.W. Smith, C.A. Ross et al., Parkinson’s disease-associated mutations in leucine-rich repeat kinase 2 augment kinase activity, Proc. Natl. Acad. Sci. U. S. A. 102 (2005), pp. 16842–16847.

[27] A.F. Kalogeropulou, E. Purlyte, F. Tonelli, S.M. Lange, M. Wightman, A.R. Prescott et al., Impact of 100 LRRK2 variants linked to Parkinson’s disease on kinase activity and microtubule binding, Biochem. J. 479 (2022), pp. 1759–1783.

[28] A.G. Henry, S. Aghamohammadzadeh, H. Samaroo, Y. Chen, K. Mou, E. Needle et al., Pathogenic LRRK2 mutations, through increased kinase activity, produce enlarged lysosomes with reduced degradative capacity and increase ATP13A2 expression, Hum. Mol. Genet. 24 (2015), pp. 6013–6028.

[29] E.D. Plowey, S.J. Cherra 3rd, Y.-J. Liu and C.T. Chu, Role of autophagy in G2019S-LRRK2-associated neurite shortening in differentiated SH-SY5Y cells, J. Neurochem. 105 (2008), pp. 1048–1056.

[30] J. Alegre-Abarrategui, H. Christian, M.M.P. Lufino, R. Mutihac, L.L. Venda, O. Ansorge et al., LRRK2 regulates autophagic activity and localizes to specific membrane microdomains in a novel human genomic reporter cellular model, Hum. Mol. Genet. 18 (2009), pp. 4022–4034.

[31] S. Matta, K. Van Kolen, R. da Cunha, G. van den Bogaart, W. Mandemakers, K. Miskiewicz et al., LRRK2 controls an EndoA phosphorylation cycle in synaptic endocytosis, Neuron 75 (2012), pp. 1008–1021.

[32] H.J. Cho, J. Yu, C. Xie, P. Rudrabhatla, X. Chen, J. Wu et al., Leucine-rich repeat kinase 2 regulates Sec16A at ER exit sites to allow ER-Golgi export, EMBO J. 33 (2014), pp. 2314–2331.

[33] R. Di Maio, E.K. Hoffman, E.M. Rocha, M.T. Keeney, L.H. Sanders, B.R. De Miranda et al., LRRK2 activation in idiopathic Parkinson’s disease, Sci. Transl. Med. 10 (2018),.

[34] K.B. Fraser, A.B. Rawlins, R.G. Clark, R.N. Alcalay, D.G. Standaert, N. Liu, et al., Ser(P)-1292 LRRK2 in urinary exosomes is elevated in idiopathic Parkinson’s disease, Mov. Disord. 31 (2016), pp. 1543–1550.

[35] S. Wang, S. Unnithan, N. Bryant, A. Chang, L.S. Rosenthal, A. Pantelyat et al., Elevated urinary Rab10 phosphorylation in idiopathic Parkinson disease, Mov. Disord. 37 (2022), pp. 1454–1464.

[36] L. Petropoulou-Vathi, A. Simitsi, P.-E. Valkimadi, M. Kedariti, L. Dimitrakopoulos, C. Koros, et al., Distinct profiles of LRRK2 activation and Rab GTPase phosphorylation in clinical samples from different PD cohorts, NPJ Parkinsons Dis. 8 (2022), pp. 73.

[37] T. Eguchi, T. Kuwahara, M. Sakurai, T. Komori, T. Fujimoto, G. Ito et al., LRRK2 and its substrate Rab GTPases are sequentially targeted onto stressed lysosomes and maintain their homeostasis, Proceedings of the National Academy of Sciences 115 (2018), pp. E9115–E9124.

[38] S. Herbst, P. Campbell, J. Harvey, E.M. Bernard, V. Papayannopoulos, N.W. Wood et al., LRRK2 activation controls the repair of damaged endomembranes in macrophages, EMBO J. 39 (2020), pp. e104494.

[39] L. Bonet-Ponce, A. Beilina, C.D. Williamson, E. Lindberg, J.H. Kluss, S. Saez-Atienzar et al., LRRK2 mediates tubulation and vesicle sorting from lysosomes, Sci. Adv. 6 (2020), pp. eabb2454.

[40] T. Kuwahara, K. Funakawa, T. Komori, M. Sakurai, G. Yoshii, T. Eguchi et al., Roles of lysosomotropic agents on LRRK2 activation and Rab10 phosphorylation, Neurobiol. Dis. 145 (2020), pp. 105081.

[41] K. Ito, M. Araki, Y. Katai, Y. Nishimura, S. Imotani, H. Inoue et al., Pathogenic LRRK2 compromises the subcellular distribution of lysosomes in a Rab12-RILPL1-dependent manner, FASEB J. 37 (2023), pp. e22930.

[42] S.E. Cason and E.L.F. Holzbaur, Axonal transport of autophagosomes is regulated by dynein activators JIP3/JIP4 and ARF/RAB GTPases, J. Cell Biol. 222 (2023),.

[43] L. Bonet-Ponce, T. Tegicho, N. Fernandez-Martinez, I.A. Rozenberg, M. Ashriem, A. Beilina et al., JIP4 and RILPL1 utilize opposing motor force to dynamically regulate lysosomal tubulation, J. Cell Biol. 224 (2025),.

[44] A. Bentley-DeSousa, A. Roczniak-Ferguson and S.M. Ferguson, A STING-CASM-GABARAP pathway activates LRRK2 at lysosomes, J. Cell Biol. 224 (2025),.

[45] B.A. Durafourt, C.S. Moore, D.A. Zammit, T.A. Johnson, F. Zaguia, M.-C. Guiot et al., Comparison of polarization properties of human adult microglia and blood-derived macrophages, Glia 60 (2012), pp. 717–727.

[46] F.O. Martinez and S. Gordon, The M1 and M2 paradigm of macrophage activation: time for reassessment, F1000Prime Rep. 6 (2014), pp. 13.

[47] R. Orihuela, C.A. McPherson and G.J. Harry, Microglial M1/M2 polarization and metabolic states: Microglia bioenergetics with acute polarization, Br. J. Pharmacol. 173 (2016), pp. 649–665.

[48] S.M. Haag, M.F. Gulen, L. Reymond, A. Gibelin, L. Abrami, A. Decout et al., Targeting STING with covalent small-molecule inhibitors, Nature 559 (2018), pp. 269–273.

[49] B. Hussain, Y. Xie, U. Jabeen, D. Lu, B. Yang, C. Wu et al., Activation of STING based on its structural features, Front. Immunol. 13 (2022), pp. 808607.

[50] S.L. Ergun, D. Fernandez, T.M. Weiss and L. Li, STING polymer structure reveals mechanisms for activation, hyperactivation, and inhibition, Cell 178 (2019), pp. 290–301.e10.

[51] P. Song, W. Yang, K.F. Lou, H. Dong, H. Zhang, B. Wang et al., UNC13D inhibits STING signaling by attenuating its oligomerization on the endoplasmic reticulum, EMBO Rep. 23 (2022), pp. e55099.

[52] R. Chan, X. Cao, S.L. Ergun, E. Njomen, S.R. Lynch, C. Ritchie et al., Human STING oligomer function is governed by palmitoylation of an evolutionarily conserved cysteine, bioRxiv, 2023,, pp. 2023.08.11.553045.

[53] A.J. Lara Ordónez, B. Fernández, E. Fdez, M. Romo-Lozano, J. Madero-Pérez, E. Lobbestael et al., RAB8, RAB10 and RILPL1 contribute to both LRRK2 kinase-mediated centrosomal cohesion and ciliogenesis deficits, Hum. Mol. Genet. 28 (2019), pp. 3552–3568.

[54] E. Fdez, J. Madero-Pérez, A.J. Lara Ordóñez, Y. Naaldijk, R. Fasiczka, A. Aiastui et al., Pathogenic LRRK2 regulates centrosome cohesion via Rab10/RILPL1-mediated CDK5RAP2 displacement, iScience 25 (2022), pp. 104476.

[55] J. Madero-Pérez, E. Fdez, B. Fernández, A.J. Lara Ordóñez, M. Blanca Ramírez, P. Gómez-Suaga et al., Parkinson disease-associated mutations in LRRK2 cause centrosomal defects via Rab8a phosphorylation, Mol. Neurodegener. 13 (2018), pp. 3.

[56] Y. Sobu, P.S. Wawro, H.S. Dhekne, W.M. Yeshaw and S.R. Pfeffer, Pathogenic LRRK2 regulates ciliation probability upstream of tau tubulin kinase 2 via Rab10 and RILPL1 proteins, Proc. Natl. Acad. Sci. U. S. A. 118 (2021), pp. e2005894118.

[57] S.S. Khan, Y. Sobu, H.S. Dhekne, F. Tonelli, K. Berndsen, D.R. Alessi et al., Pathogenic LRRK2 control of primary cilia and Hedgehog signaling in neurons and astrocytes of mouse brain, Elife 10 (2021),.

[58] X. Li, H. Zhu, B.T. Huang, X. Li, H. Kim, H. Tan et al., RAB12-LRRK2 complex suppresses primary ciliogenesis and regulates centrosome homeostasis in astrocytes, Nat. Commun. 15 (2024), pp. 8434.

[59] C.M. Ott, R. Torres, T.-S. Kuan, A. Kuan, J. Buchanan, L. Elabbady et al., Ultrastructural differences impact cilia shape and external exposure across cell classes in the visual cortex, Curr. Biol. 34 (2024), pp. 2418–2433.e4.

[60] P. Volos, K. Fujise and N.M. Rafiq, Roles for primary cilia in synapses and neurological disorders, Trends Cell Biol. 35 (2025), pp. 6–10.

[61] R. Ulferts, E. Marcassa, L. Timimi, L.C. Lee, A. Daley, B. Montaner et al., Subtractive CRISPR screen identifies the ATG16L1/vacuolar ATPase axis as required for non-canonical LC3 lipidation, Cell Rep. 37 (2021), pp. 109899.

[62] Y. Tanaka and Z.J. Chen, STING specifies IRF3 phosphorylation by TBK1 in the cytosolic DNA signaling pathway, Sci. Signal. 5 (2012), pp. ra20.

[63] T. Saitoh, N. Fujita, T. Hayashi, K. Takahara, T. Satoh, H. Lee et al., Atg9a controls dsDNA-driven dynamic translocation of STING and the innate immune response, Proc. Natl. Acad. Sci. U. S. A. 106 (2009), pp. 20842–20846.

[64] Y. Kuchitsu, K. Mukai, R. Uematsu, Y. Takaada, A. Shinojima, R. Shindo et al., STING signalling is terminated through ESCRT-dependent microautophagy of vesicles originating from recycling endosomes, Nat. Cell Biol. 25 (2023), pp. 453–466.

[65] Y. Kuchitsu and T. Taguchi, STINGing organelle surface with acid, EMBO Rep. 25 (2024), pp. 1708–1710.

[66] H. Kemmoku, K. Takahashi, K. Mukai, T. Mori, K.M. Hirosawa, F. Kiku et al., Single-molecule localization microscopy reveals STING clustering at the trans-Golgi network through palmitoylation-dependent accumulation of cholesterol, Nat. Commun. 15 (2024), pp. 220.

[67] T. Kuwahara and T. Iwatsubo, CASM mediates LRRK2 recruitment and activation under lysosomal stress, Autophagy 20 (2024), pp. 1692–1693.

[68] T. Eguchi, M. Sakurai, Y. Wang, C. Saito, G. Yoshii, T. Wileman et al., The V-ATPase-ATG16L1 axis recruits LRRK2 to facilitate the lysosomal stress response, J. Cell Biol. 223 (2024),.

[69] Y. Lei and D.J. Klionsky, The coordination of V-ATPase and ATG16L1 is part of a common mechanism of non-canonical autophagy, Autophagy 18 (2022), pp. 2267–2269.

[70] T. Kuwahara, G. Yoshii, M. Sakurai, S. Suenaga, H. Nakanishi, M. Jefferson et al., LRRK2 is activated by phosphatidylinositol 3-phosphate in conjunction with CASM, bioRxiv, 2025,, pp. 2025.12.16.694573.

[71] M.J. Fell, C. Mirescu, K. Basu, B. Cheewatrakoolpong, D.E. DeMong, J.M. Ellis et al., MLi-2, a potent, selective, and centrally active compound for exploring the therapeutic potential and safety of LRRK2 kinase inhibition, J. Pharmacol. Exp. Ther. 355 (2015), pp. 397–409.

[72] M. Sakurai, T. Kuwahara, S. Suenaga, S. Takatori, T. Tomita, T. Shalit, et al., Subtype*-specific secretion of extracellular vesicles by LRRK2 and Rab GTPases under lysosomal stress*, iScience 29 (2026), pp. 117203.

[73] K.F. O’Connell, Centrosomes: An acentriolar MTOC at the ciliary base, Current biology: CB, 31 (2021), pp. R730–R733.

[74] H. Kobayashi, K. Etoh, N. Ohbayashi and M. Fukuda, Rab35 promotes the recruitment of Rab8, Rab13 and Rab36 to recycling endosomes through MICAL-L1 during neurite outgrowth, Biol. Open 3 (2014), pp. 803–814.

[75] Y. Messaoud-Nacer, E. Culerier, S. Rose, I. Maillet, N. Rouxel, S. Briault et al., STING agonist diABZI induces PANoptosis and DNA mediated acute respiratory distress syndrome (ARDS), Cell Death Dis. 13 (2022), pp. 269.

[76] T. Taniguchi and A. Takaoka, Type I interferon system and IRF family of transcription factors in host defense regulation, Proc. Jpn. Acad. Ser. B Phys. Biol. Sci. 81 (2005), pp. 1–13.

[77] F. Ma, B. Li, Y. Yu, S.S. Iyer, M. Sun and G. Cheng, Positive feedback regulation of type I interferon by the interferon-stimulated gene STING, EMBO Rep. 16 (2015), pp. 202– 212.

[78] S. Yum, M. Li, Y. Fang and Z.J. Chen, TBK1 recruitment to STING activates both IRF3 and NF-κB that mediate immune defense against tumors and viral infections, Proc. Natl. Acad. Sci. U. S. A. 118 (2021), pp. e2100225118.

[79] P.J. Tapia, J.A. Martina, P.S. Contreras, A. Prashar, E. Jeong, D. De Nardo et al., TFEB and TFE3 regulate STING1-dependent immune responses by controlling type I interferon signaling, Autophagy 21 (2025), pp. 2028–2045.

[80] B. Lv, W.A. Dion, H. Yang, J. Xun, D.-H. Kim, B. Zhu et al., A TBK1-independent primordial function of STING in lysosomal biogenesis, Mol. Cell 84 (2024), pp. 3979–3996.e9.

[81] Y. Xu, Q. Wang, J. Wang, C. Qian, Y. Wang, S. Lu et al., The cGAS-STING pathway activates transcription factor TFEB to stimulate lysosome biogenesis and pathogen clearance, Immunity 58 (2025), pp. 309–325.e6.

[82] N. Yadavalli and S.M. Ferguson, LRRK2 suppresses lysosome degradative activity in macrophages and microglia through MiT-TFE transcription factor inhibition, Proc. Natl. Acad. Sci. U. S. A. 120 (2023), pp. e2303789120.

[83] M. Blanca Ramírez, A.J. Lara Ordóñez, E. Fdez, J. Madero-Pérez, A. Gonnelli, M. Drouyer et al., GTP binding regulates cellular localization of Parkinson’s disease-associated LRRK2, Hum. Mol. Genet. 26 (2017), pp. 2747–2767.

[84] J. Madero-Pérez, B. Fernández, A.J. Lara Ordóñez, E. Fdez, E. Lobbestael, V. Baekelandt et al., RAB7L1-mediated relocalization of LRRK2 to the Golgi complex causes centrosomal deficits via RAB8A, Front. Mol. Neurosci. 11 (2018), pp. 417.

[85] Z. Gao, B. Wang and L. Zhang, Activated STING: an ion channel to trigger non-interferon-related functions, Signal Transduction and Targeted Therapy, 8 (2023), pp. 388.

[86] D.E. Johnson, P. Ostrowski, V. Jaumouillé and S. Grinstein, The position of lysosomes within the cell determines their luminal pH, J. Cell Biol. 212 (2016), pp. 677–692.

[87] Y. Sasazawa, S. Souma, N. Furuya, Y. Miura, S. Kazuno, S. Kakuta et al., Oxidative stress-induced phosphorylation of JIP4 regulates lysosomal positioning in coordination with TRPML1 and ALG2, EMBO J. 41 (2022), pp. e111476.

[88] G. Villari, C. Enrico Bena, M. Del Giudice, N. Gioelli, C. Sandri, C. Camillo et al., Distinct retrograde microtubule motor sets drive early and late endosome transport, EMBO J. 39 (2020), pp. e103661.

[89] O. Ullrich, S. Reinsch, S. Urbé, M. Zerial and R.G. Parton, Rab11 regulates recycling through the pericentriolar recycling endosome, J. Cell Biol. 135 (1996), pp. 913–924.

[90] J.H. Kluss, A. Beilina, C.D. Williamson, P.A. Lewis, M.R. Cookson and L. Bonet-Ponce, Lysosomal positioning regulates Rab10 phosphorylation at LRRK2+ lysosomes, Proc. Natl. Acad. Sci. U. S. A. 119 (2022), pp. e2205492119.

[91] T. Malankhanova, Z. Liu, E. Xu, N. Bryant, K.W. Sung, H. Li et al., LRRK2 interactions with microtubules are independent of LRRK2-mediated Rab phosphorylation, EMBO Rep. 26 (2025), pp. 3445–3466.

[92] H.S. Dhekne, I. Yanatori, R.C. Gomez, F. Tonelli, F. Diez, B. Schüle et al., A pathway for Parkinson’s Disease LRRK2 kinase to block primary cilia and Sonic hedgehog signaling in the brain, Elife 7 (2018),.

[93] A. Iguchi, S. Takatori, S. Kimura, H. Muneto, K. Wang, H. Etani, et al., INPP5D modulates TREM2 loss-of-function phenotypes in a β-amyloidosis mouse model, iScience 26 (2023), pp. 106375.

[94] T. Abe, T. Kuwahara, S. Suenaga, M. Sakurai, S. Takatori and T. Iwatsubo, Lysosomal stress drives the release of pathogenic α-synuclein from macrophage lineage cells via the LRRK2-Rab10 pathway, iScience 27 (2024), pp. 108893.

