## Supplemental Figures for "STING activation drives pericentrosomal positioning of acidic organelles via the VAIL-LRRK2-Rab35 pathway"

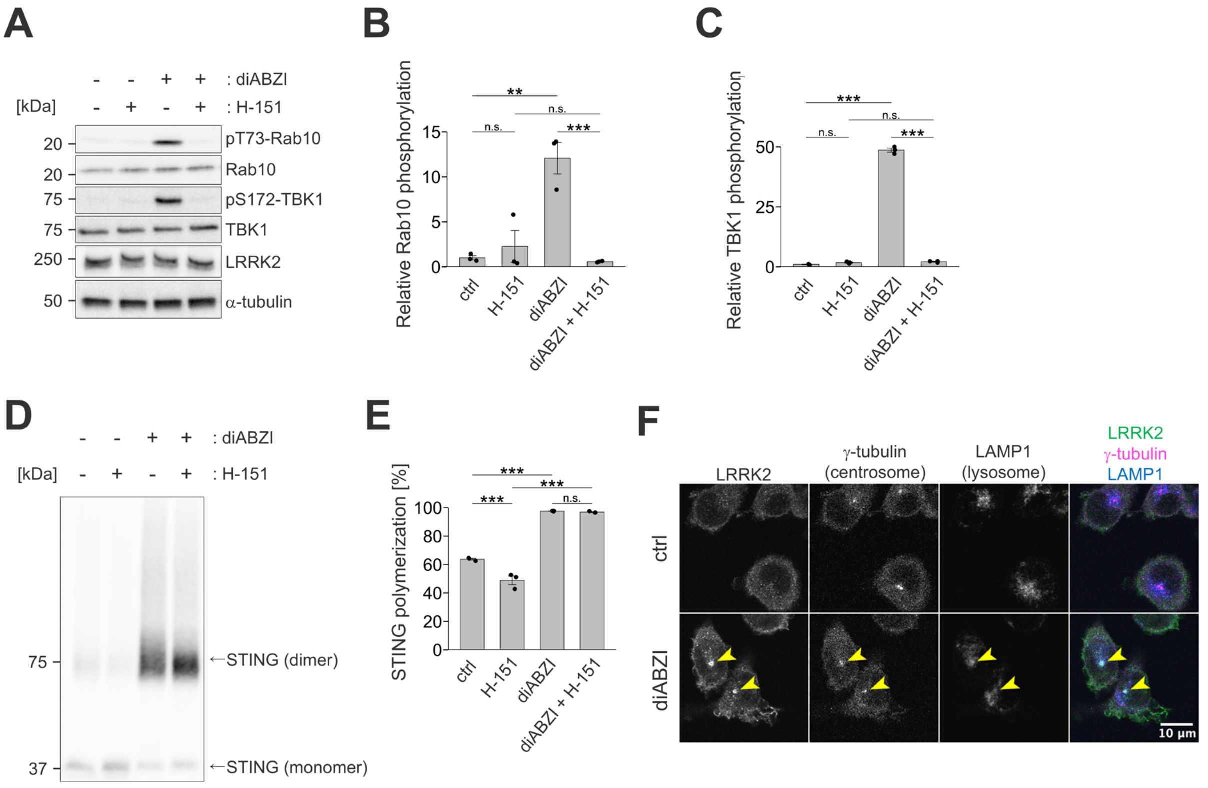


**Figure S1.** A STING agonist diABZI also induces LRRK2 activation and translocation.

(**A**) Immunoblotting of lysates of RAW264.7 cells primed by IFNγ and treated with diABZI and/or STING inhibitor H-151 for two hours. Representative images from three independent sample sets. (**B**, **C**) Quantification of Rab10 phosphorylation (B) and TBK1 phosphorylation (C). Statistical significance was assessed by two-way ANOVA with Tukey's post-hoc test. (**D**) Non-reducing immunoblotting of the same sets of cell lysates as in A probed with an anti-STING antibody. (**E**) Quantification of STING dimerization. Statistical significance was assessed by two-tailed Welch's *t*-tests with Holm-Bonferroni's correction. (**F**) Immunocytochemical analysis of IFNγ-primed MG6 cells treated with diABZI. **: $p < 0.01$, ***: $p < 0.001$.

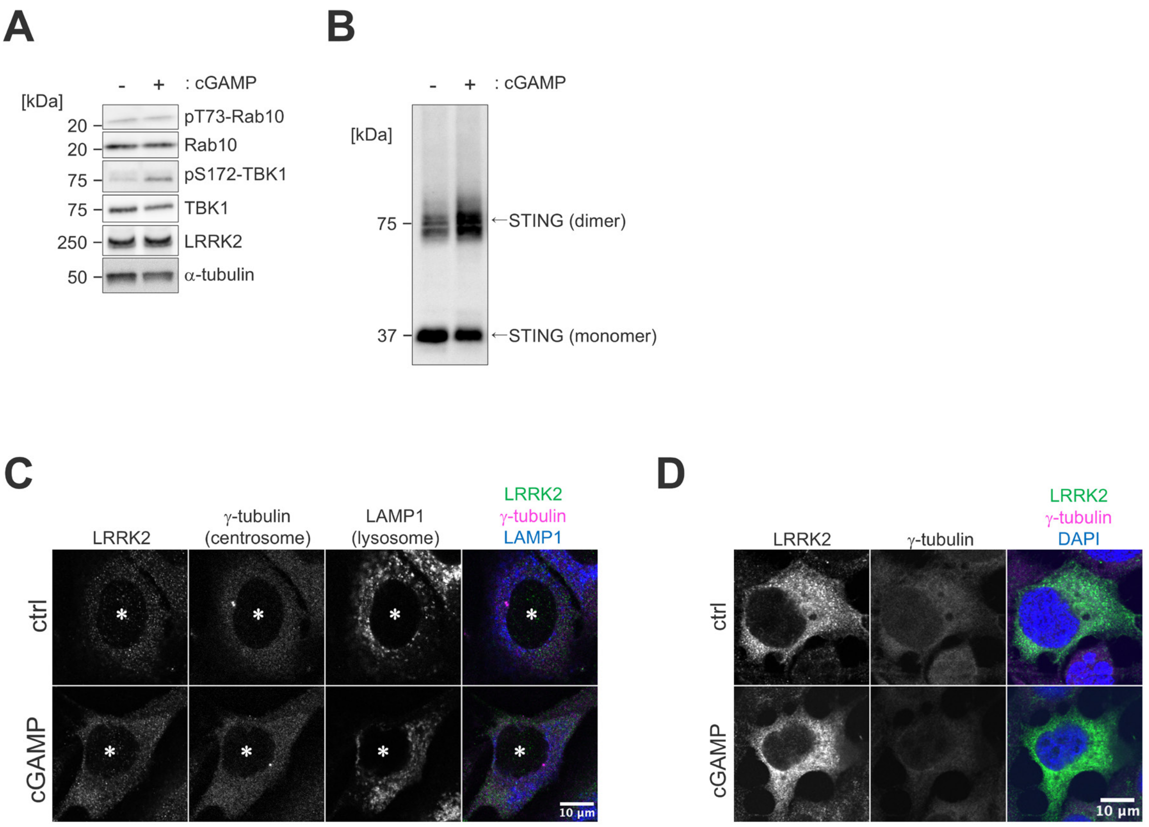


**Figure S2.** Cell type specificity in STING-mediated translocation of LRRK2 to the pericentrosomal area.

(**A**) Immunoblotting of lysates of NIH/3T3 cells treated with or without cGAMP for six hours. Representative images from two independent sample sets. (**B**) Non-reducing immunoblotting of the same sets of cell lysates as in A probed with an anti-STING antibody. (**C**) Immunocytochemical analysis of NIH/3T3 cells treated with or without cGAMP. (**D**) Immunocytochemical analysis of HEK293 cells transfected to express 3×FLAG-LRRK2 transiently and treated with cGAMP.

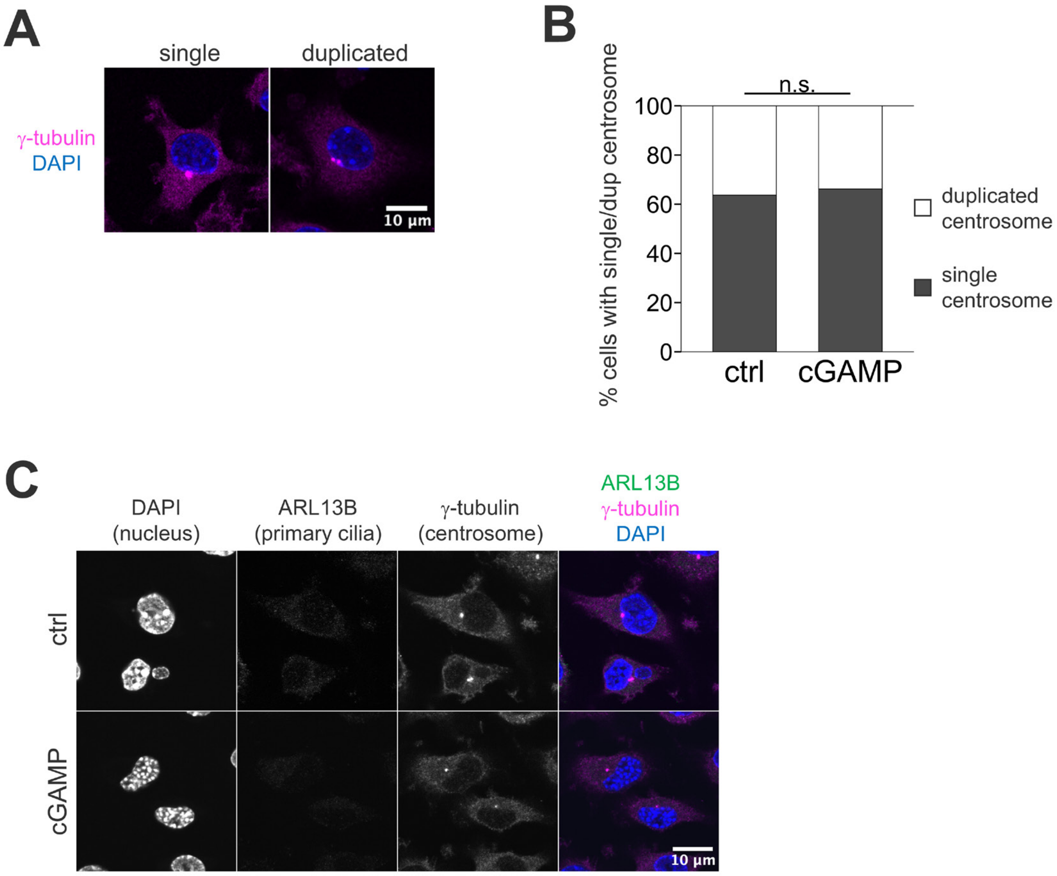


**Figure S3.** Centrosomal cohesion and primary cilia are not affected upon STING activation. (**A**) Single or duplicated (split) centrosomes in MG6 cells, stained with an anti-γ-tubulin antibody. (**B**) The ratio of the split centrosomes in MG6 cells treated with or without cGAMP for six hours. Statistical significance was assessed by one-way ANOVA. n.s.: not significant (**C**) Immunocytochemical analysis of ARL13b, a marker protein for primary cilia, in MG6 cells.

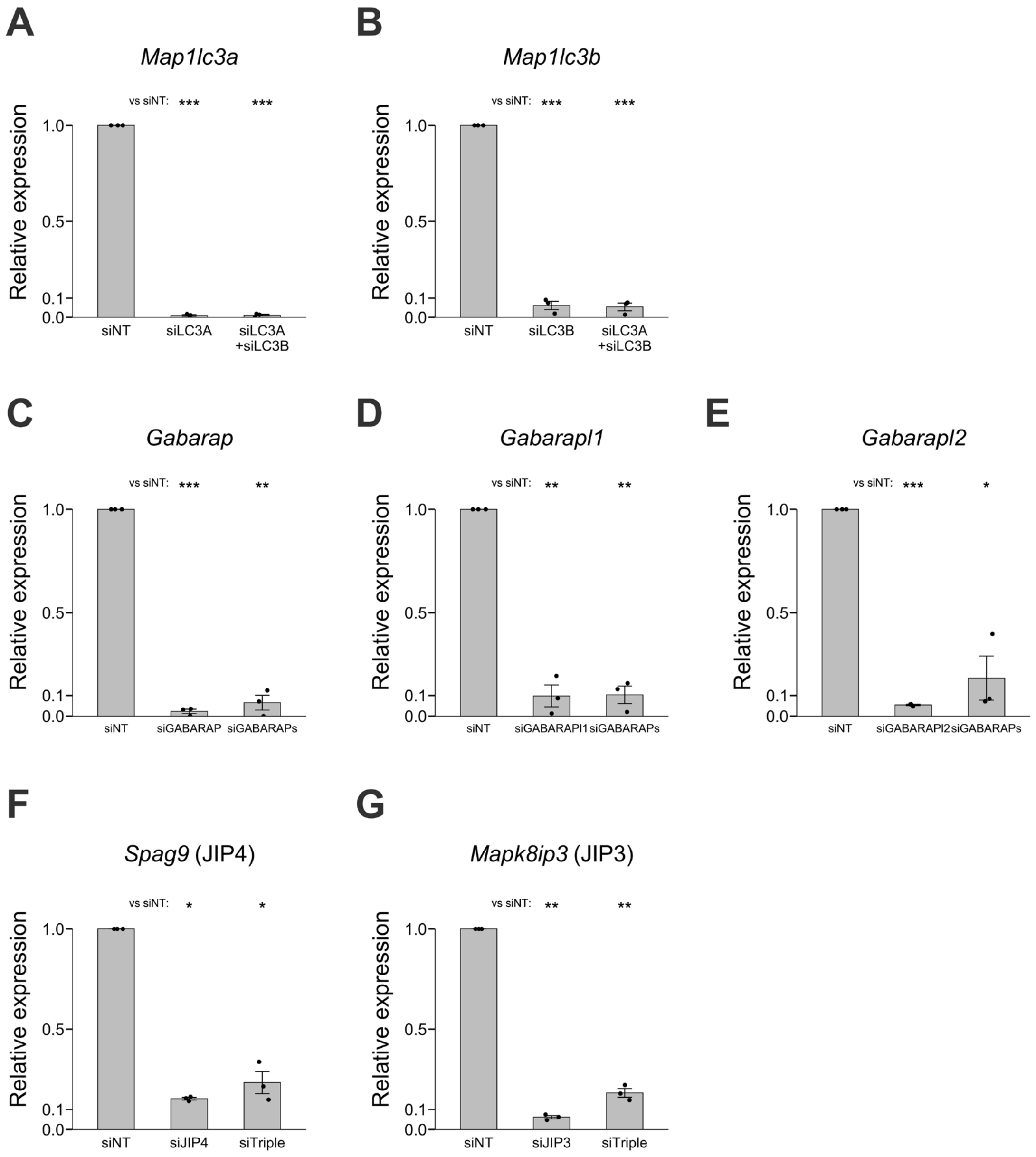


**Figure S4.** Confirmation of the knockdown efficiencies in RAW264.7 cells.
(**A-E**) qPCR analysis of transcripts of each ATG8 gene to confirm the knockdown efficiency in each condition. Statistical significance was assessed by paired two-tailed *t*-tests on ΔCt values (siTarget vs siNT within each biological replicate), with Holm correction for multiple comparisons. (**F**, **G**) qPCR analysis for transcripts of each kinesin/dynein adaptor gene to confirm the knockdown efficiency in each condition. Statistical significance was assessed by paired two-tailed *t*-tests on ΔCt values. *: $p < 0.05$, **: $p < 0.01$.

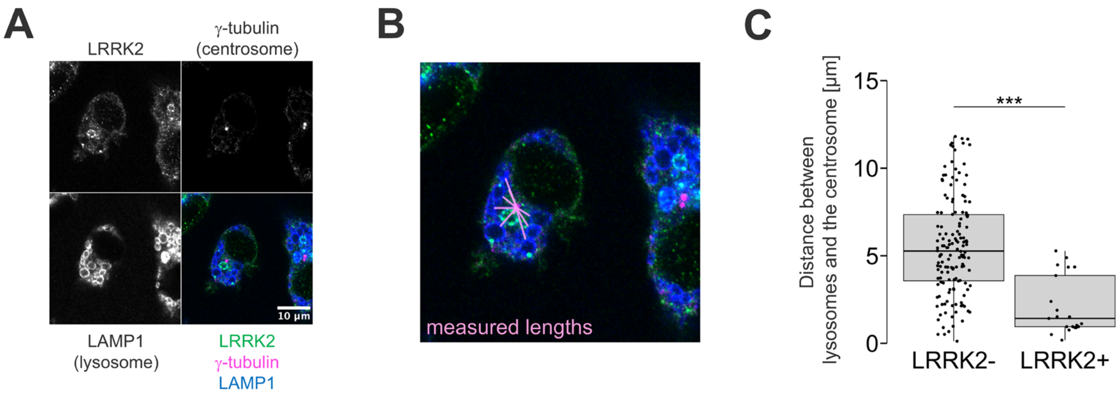


**Figure S5.** CQ treatment induces recruitment of LRRK2 to lysosomes near the centrosome. (**A**) Immunocytochemical analysis of RAW264.7 cells treated with chloroquine for three hours and stained for LRRK2, γ-tubulin, and LAMP1. (**B**) Representative measuring of distances from the centrosome to enlarged lysosomes. (**C**) Distances between lysosomes and the centrosome, measured for LRRK2-negative lysosomes and LRRK2-positive ones. Statistical significance was assessed by one-way ANOVA. 150 or 21 lysosomes were included in each group. ***: $p < 0.001$.

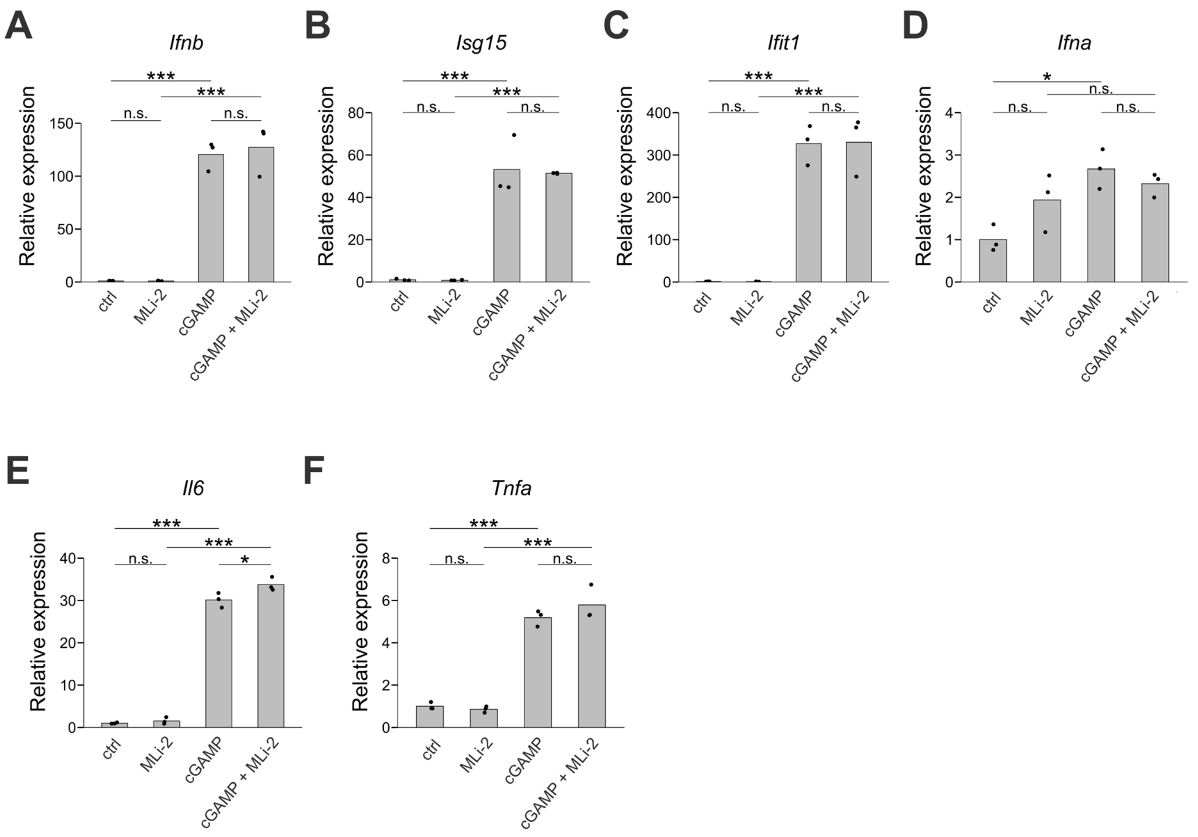


**Figure S6.** LRRK2 kinase activity does not modify the STING immune pathways.

(**A**-**F**) mRNAs from RAW264.7 cells primed by IFNγ and treated with cGAMP and/or MLi-2 for six hours were quantified by qPCR for the indicated interferon-related or inflammatory genes. Statistical significance was assessed by two-way ANOVA with Tukey's post-hoc tests. n.s.: not significant, *: $p < 0.05$, ***: $p < 0.001$.

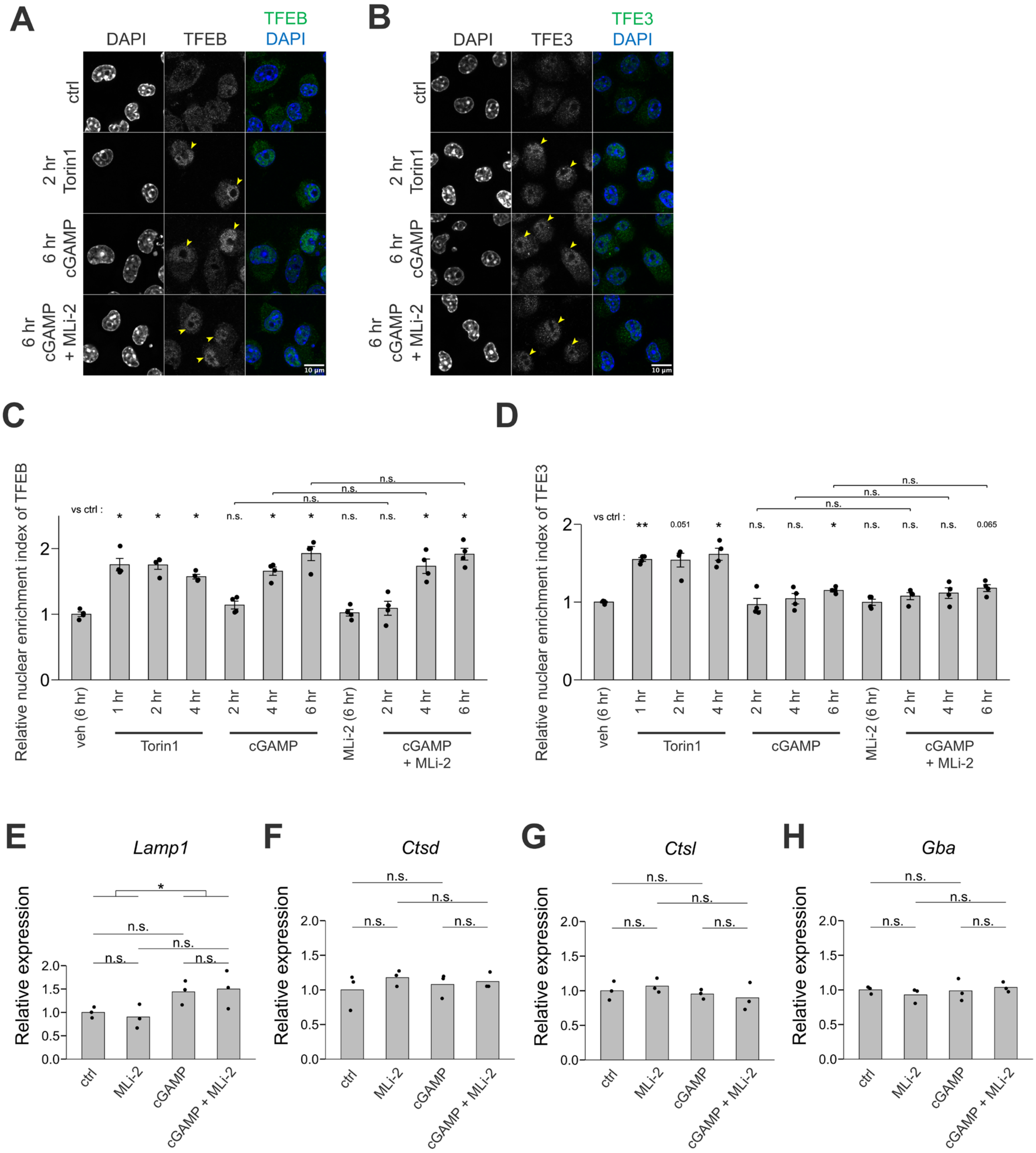


**Figure S7.** LRRK2 kinase activity does not modify the STING-VAIL-TFE pathways. (**A**, **B**) Immunocytochemical analysis for TFEB (A) and TFE3 (B) for MG6 cells primed by IFNγ and treated with Torin1 or cGAMP and/or MLi-2. (**C**, **D**) Signal enrichment in nuclei for TFEB (C) and TFE3 (D) were quantified for each observation and averaged. Data from four observations were included. Thirteen *t*-tests were conducted and *p*-values adjusted by Holm-Bonferroni's correction are shown. (**E-H**) mRNAs from RAW264.7 cells primed by IFNγ and treated with cGAMP and/or MLi-2 for six hours were quantified by qPCR for the indicated lysosomal genes. Statistical significance was assessed by two-way ANOVA with Tukey's post-hoc tests. n.s.: not significant, *: $p < 0.05$.

| index | target | catalog id |
|---|---|---|
| 1 | *Atg16l1* | D-051699-01 |
| 2 | *Map1lc3a* | M-056203-00 |
| 3 | *Map1lc3b* | D-040989-02 |
| 4 | *Gabarap* | D-041776-02 |
| 5 | *Gabarapl1* | D-040444-01 |
| 6 | *Gabarapl2* | D-059605-02 |
| 7 | *Rab8a* | M-040860-00 |
| 8 | *Rab8b* | M-055301-00 |
| 9 | *Rab10* | M-040862-01 |
| 10 | *Rab12* | M-040865-01 |
| 11 | *Rab35* | D-042604-00 |
| 12 | *Rilpl1* | D-063225-03 |
| 13 | *Spag9* | D-048195-04 |
| 14 | *Mapk8ip3* | D-043334-04 |

**Table S1.** siRNAs used in this study.

| index | name | sequence (5'->3') |
|---|---|---|
| 1 | Gapdh_Fw | CTGCAGCCTCGTCCCGTA |
| 2 | Gapdh_Rv | GTGACCAGGCGCCCAATAC |
| 3 | Atg16l1_Fw | TCAGCCGAGGGTTCTCTTTA |
| 4 | Atg16l1_Rv | GCTCTGCTTCCTTTGTCCAC |
| 5 | Map1lc3a_Fw | CTGTAAGGAGGTGCAGCAGA |
| 6 | Map1lc3a_Rv | GTCTGGGACCAGAAACTTGG |
| 7 | Map1lc3b_Fw | CCGTCCGAGAAGACCTTCAA |
| 8 | Map1lc3b_Rv | TCGCTCTATAATCACTGGGATCTTG |
| 9 | Gabarap_Fw | ACAATGTCATTCCACCCACCA |
| 10 | Gabarap_Rv | TCAGGTACAGCAGCTTCACAG |
| 11 | Gabarapl1_Fw | AGGACCACCCCTTCGAGTATC |
| 12 | Gabarapl1_Rv | GAGGGCACAAGGTACTTCCTC |
| 13 | Gabarapl2_Fw | GAAGATCAGAGCGAAGTACCCC |
| 14 | Gabarapl2_Rv | ATCCACATGAACTGAGCCACA |
| 15 | Spag9_Fw | TGGATGAAGGAGCGGATTTACT |
| 16 | Spag9_Rv | CATTCTTCACTACATTCAGAGCGT |
| 17 | Mapk8ip3_Fw | CCTGAAACCCGTCTGGAGC |
| 18 | Mapk8ip3_Rv | AAATCATCGCGCACTGAAAAC |
| 19 | Ifnb_Fw | GGCGGACTTCAAGATCCCTA |
| 20 | Ifnb_Rv | AGTCTCATTCCACCCAGTGC |
| 21 | Isg15_Fw | GGGGCCACAGCAACATCTAT |
| 22 | Isg15_Rv | GGCTTTAGGCCATACTCCCC |
| 23 | Ifit1_Fw | TCGCGTAGACAAAGCTCTTCA |
| 24 | Ifit1_Rv | GCCTGTTTCGGGATGTCCTC |
| 25 | pan_Ifna_Fw | TGCCCAGCAGATCAAGAAGG |
| 26 | pan_Ifna_Rv | TCAGGGGAAATTCCTGCACC |
| 27 | Il6_Fw | CCGGAGAGGAGACTTCACAG |
| 28 | Il6_Rv | TCCACGATTTCCCAGAGAAC |
| 29 | Tnf_Fw | ACGGCATGGATCTCAAAGAC |
| 30 | Tnf_Rv | GTGGGTGAGGAGCACGTAGT |
| 31 | Lamp1_Fw | CTGCACACAGGATGGACCTT |
| 32 | Lamp1_Rv | CCAAACTGCAATTCCAGGGC |
| 33 | Ctsd_Fw | CTATAAGCCGGCGACCTCTG |
| 34 | Ctsd_Rv | TGAACTTGCGCAGAGGGATT |
| 35 | Ctsl_Fw | TCGACCATGGGGTTCTGTTG |
| 36 | Ctsl_Rv | GTTGTCCCGGTCTTTGGCTA |
| 37 | Gba_Fw | CCTACAGCAGGGCTCTTCAC |
| 38 | Gba_Rv | CAAGCGTTGGTCATCTAGCA |

**Table S2.** Sequences of primers used for RT-qPCR in this study.
